# The Effect of Oblique Image Acquisition on the Accuracy of Quantitative Susceptibility Mapping and a Robust Tilt Correction Method

**DOI:** 10.1101/2021.11.30.470544

**Authors:** Oliver C. Kiersnowski, Anita Karsa, Stephen J. Wastling, John S. Thornton, Karin Shmueli

## Abstract

**Purpose:** Quantitative susceptibility mapping (QSM) is increasingly used for clinical research where oblique image acquisition is commonplace but its effects on QSM accuracy are not well understood.

**Theory and Methods:** The QSM processing pipeline involves defining the unit magnetic dipole kernel, which requires knowledge of the direction of the main magnetic field 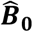 with respect to the acquired image volume axes. The direction of 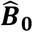 is dependent upon the axis and angle of rotation in oblique acquisition. Using both a numerical brain phantom and in-vivo acquisitions in five healthy volunteers, we analysed the effects of oblique acquisition on magnetic susceptibility maps. We compared three tilt correction schemes at each step in the QSM pipeline: phase unwrapping, background field removal and susceptibility calculation, using the root-mean-squared error and QSM-tuned structural similarity index (XSIM).

**Results:** Rotation of wrapped phase images gave severe artefacts. Background field removal with PDF gave the most accurate susceptibilities when the field map was first rotated into alignment with 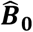. LBV and V-SHARP background field removal methods gave accurate results without tilt correction. For susceptibility calculation, thresholded *k*-space division, iterative Tikhonov regularisation and weighted linear total variation regularisation all performed most accurately when local field maps were rotated into alignment with 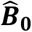 before susceptibility calculation.

**Conclusion:** For accurate QSM, oblique acquisition must be taken into account. Rotation of images into alignment with 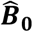 should be carried out after phase unwrapping and before background field removal. We provide open-source tilt-correction code to incorporate easily into existing pipelines: https://github.com/o-snow/QSM_TiltCorrection.git.

## Introduction

The acquisition of oblique image slices, or an oblique slab or volume in 3D MRI, is common in clinical practice to facilitate radiological viewing of brain MRI. For example, axial slices are often aligned along the subcallosal line for longitudinal studies that require consistent repositioning of acquired images^1^. Alternatively, slices may be aligned perpendicular to the principle axis of the hippocampus for accurate hippocampal volume measurements and sharper hippocampal boundary delineation^2^. Oblique slices are also acquired to reduce image artefacts from, for example, eye motion resulting in localised blurring around the eyes and ghosting along the phase encode direction^3^. Note that acquiring oblique slices does not require the subject to rotate their head, as only the acquisition volume is tilted.

Quantitative susceptibility mapping^4–6^ utilises the information in the (conventionally discarded) phase component, ϕ(**r**), of the complex MRI signal from a gradient echo (GRE) sequence to calculate the tissue magnetic susceptibility, χ. A typical QSM pipeline includes three key steps: 1) phase unwrapping of wraps present due to ϕ(**r**) being constrained to the [–π,π) interval, 2) background field removal separating the local field perturbations due to internal χ sources inside the volume of interest (e.g. the brain), Δ*B_int_*(***r***), from unwanted background field perturbations due to external sources, Δ*B_ext_*(***r***), and 3) a local-field-to-χ(**r**) calculation to solve an ill-posed inverse problem:

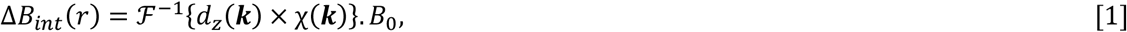

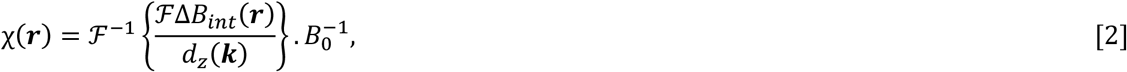

where 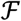 is the Fourier transform, 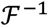 its inverse, *B*_0_ the magnetic field in Tesla, and *d_z_*(***k***) the *z*-component of the magnetic dipole in *k*-space 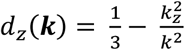 (see Equation 5).

Calculation of *d_z_*(***k***) requires knowledge of the ‘***z***’ direction of the main magnetic field, **B_0_**, with respect to the image volume acquired. Therefore, oblique acquisition must be taken into account within the QSM pipeline otherwise incorrect χ estimates arise, as suggested by a preliminary study^7^ and our preliminary data^8^. With the increase in clinical applications of QSM^9,10^, accuracy in χ estimates for oblique acquisition, typical in clinical protocols, is of paramount importance in ensuring smooth translation of QSM into clinical practice. However, accurate QSM accounting for oblique acquisition is non-trivial and there are a number of techniques proposed to account for oblique acquisition in QSM^n–14^ including the most common methods of rotating the *k*-space dipole or the image volume into alignment with ***B*_0_**. The effect of these proposed tilt correction techniques on susceptibility values has not been evaluated and it is not known at which point in the QSM pipeline these techniques should be applied. Furthermore, it is not clear what is the optimal method for taking oblique acquisition into account in the QSM pipeline: simply defining the dipole at an angle (see *DipK* or *DipIm* below) has been shown to be non-optimal^7^. Therefore, the research presented here is the first quantitative and comparative evaluation of correction methods for oblique acquisition in QSM. We used a numerical phantom to carry out a comprehensive analysis of the effect of oblique acquisition on each step of the QSM pipeline, and propose three tilt correction schemes, analysing their effects on susceptibility values when applied at different points in the QSM pipeline. We also acquired several images, in five healthy volunteers, with volumes tilted at different angles and performed the same analysis of the effects of tilting and correction schemes in vivo. We provide open-source tilt-correction code at https://github.com/o-snow/QSM_TiltCorrection.git that uses the header information from NIfTI^15^ format images to correctly orient image volumes and account for tilted acquisition for accurate QSM.

## Theory

To accurately model the magnetic dipole kernel required for the field-to-χ calculation and, in some cases, for background field removal, it is necessary to know where the magnetic field ***B*_0_** lies in acquired MRI images. Defining the two coordinate systems of interest^7^ as the acquired image frame 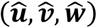 and the scanner frame 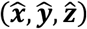, the main magnetic field can be written as ***B***_0,*im*_ and ***B***_0*sc*_ in the image and scanner frames, respectively:

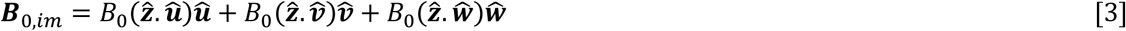

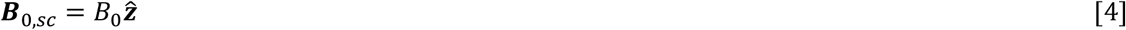

In the case of non-oblique acquisition, the coordinate systems are aligned and 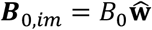 in the image frame (Figure 1, left).

**Figure 1:**
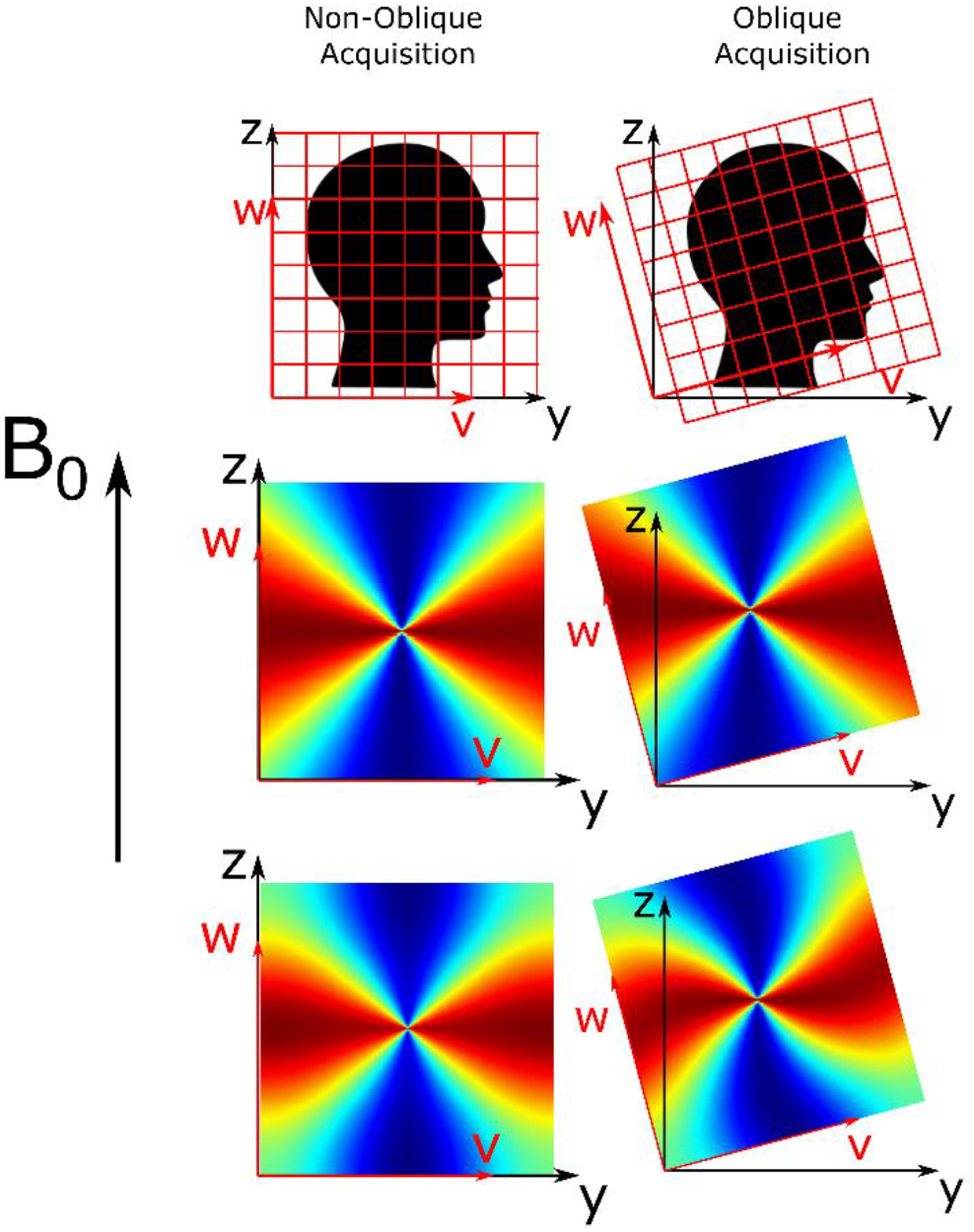
Non-oblique and oblique acquisition about the ***x***-axis (***u***-axis) of axial slices (top row) with corresponding *k*-space dipoles (middle row) and image-space dipoles (bottom row). The image axes (***w, v, w***) and scanner axes (***x, y, z***) are shown in red and black, respectively. Note that the rotation axis is at the centre of the image.

For the local field, Δ*B_int_*(***r***), to *χ*(***r***) calculation (Equation 2), the magnetic dipole kernel must be calculated. Throughout this paper, references will be made to the dimensionless *k*-space dipole, *d_z_*(***k***) (Figure 1, middle row), and the dimensionless ‘image-space dipole’ defined in image space and Fourier transformed into k-space, *d_z,im_*(***k***) (Figure 1, bottom row), kernels defined as follows^16,17^:

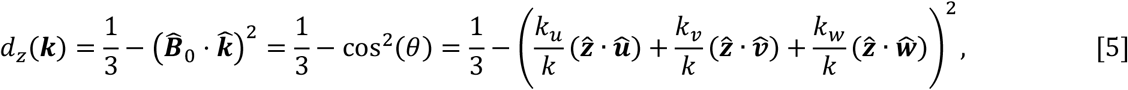

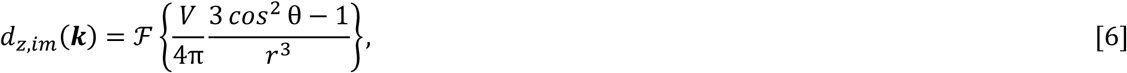

with 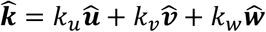 the unit vector of *k*, 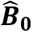 is the unit vector of ***B*_0_**, ***V*** is the voxel volume, **θ** is the angle between 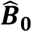 and 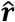, the unit vector of ***r*** in image space, where 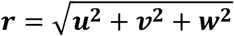, and 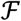 is the Fourier transform. The periodicity of the Fourier transform constrains the boundaries of *k*-space, resulting in the dipole pattern becoming fixed along those boundaries. This causes a rotated image-space dipole to appear twisted, sheared or distorted (Figure 1, bottom row). It is possible to obtain the direction of 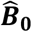 relative to the (tilted) image axes from the image headers (e,g. DICOM or Nifti format) and, therefore, to correctly calculate the magnetic dipole kernel using either equation 5 or 6.

## Methods

To determine the optimal method for taking oblique acquisition into account in the QSM pipeline we investigated three proposed tilt correction schemes, and, for comparison, an uncorrected analysis pipeline (Figure 2):

1. ***RotPrior***: rotation of the oblique image into alignment with the scanner frame prior to phase unwrapping, background field removal or the susceptibility calculation method. In this method, the dipole is defined in *k*-space in the scanner frame (using equation 5)
2. ***DipK***: the image is left unaligned to the scanner frame and the dipole used is defined in *k-* space in the oblique image frame (using equation 5). This is the default tilt correction method implemented in popular QSM toolboxes^14,18,19^. However, this method often requires the user to input the corrected ***B*_0_** direction, which is optional in many of these toolboxes.
3. ***DipIm***: the image is left unaligned to the scanner frame and the dipole used is defined in image-space in the oblique image frame (using equation 6).
4. ***NoRot***: the oblique image is left unaligned to the scanner frame, and the *k*-space dipole is mistakenly defined in the scanner frame (using equation 4) and is thereby misaligned to the true magnetic field direction, ***B*_0_**. This is the uncorrected method which can easily result from users failing to input (the correct) ***B*_0_** direction.

**Figure 2:**
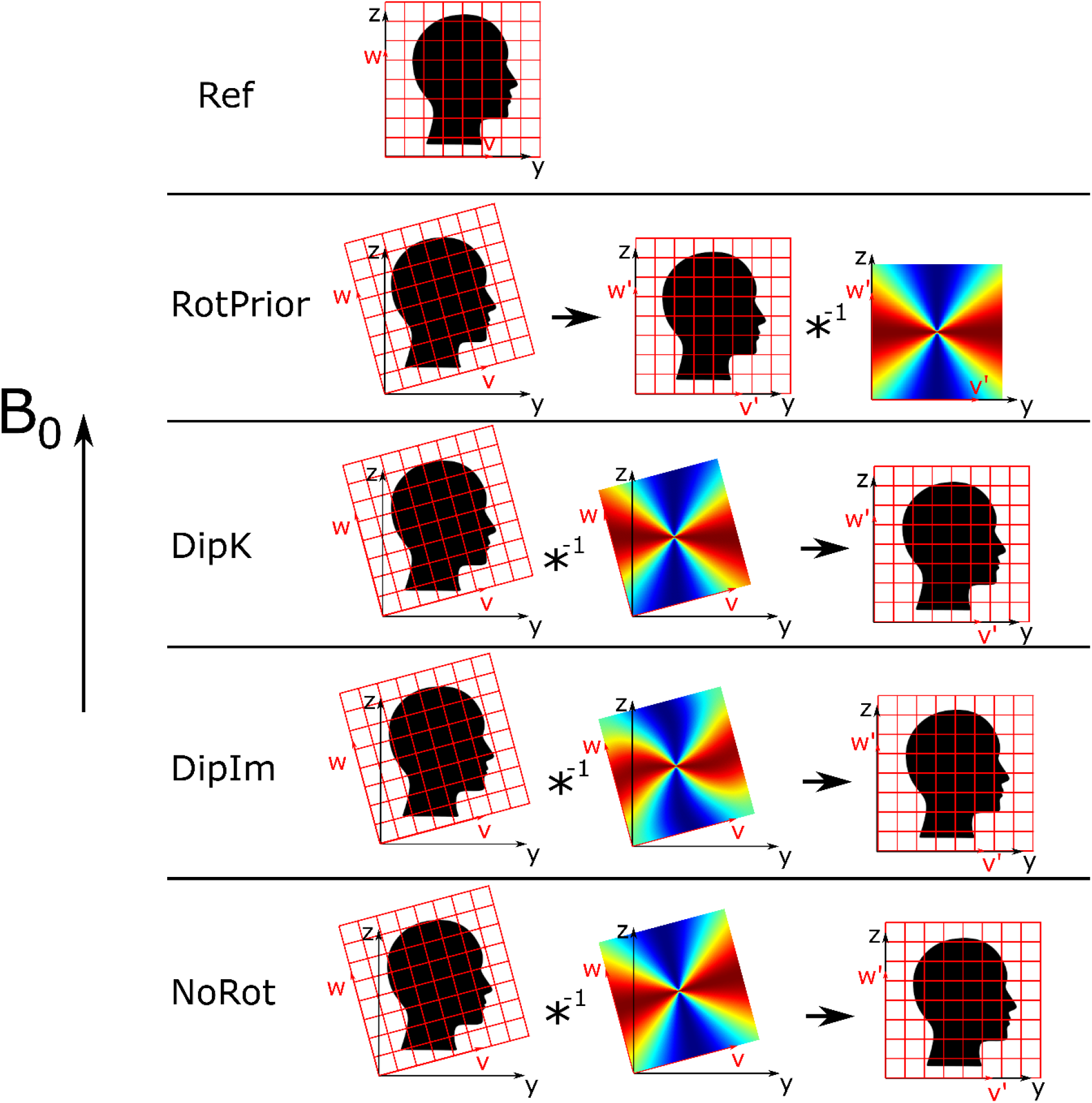
All tilt correction schemes including the reference, non-oblique acquisition for rotations about the *x*-axis. The native (oblique) image space 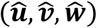 was transformed to 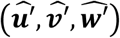 aligned with the scanner frame. The black arrow denotes rotation into the scanner frame of reference. *DipK, DipIm* and *NoRot* were rotated back into the reference (scanner) frame post-correction to facilitate comparisons. *RotPrior* and *NoRot* still apply when no dipole is used.

These schemes are the general names of the methods that we applied at different points in the pipeline (i.e. before phase unwrapping, background field removal, or susceptibility calculation) and for different methods or algorithms. For example, as no dipole kernel is necessary for phase unwrapping (and we substitute the dipole operations illustrated in Figure 2 with 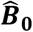-orientation-independent unwrapping operations), we have called the only two schemes appropriate before unwrapping *RotPrior* and *NoRot* where the image volume is rotated prior to unwrapping and after, respectively.

All rotations were carried out about the *x*-axis (*u*-axis) to simulate single oblique acquisition, the *y*-axis (*v*-axis) for confirmation, and about the *y = x* axis (*v = u* axis) to simulate double oblique acquisition. Rotations were undertaken using FSL FLIRT^20^ with trilinear interpolation. To facilitate comparisons, all images left in the image-frame after correction (*DipK, DipIm* and *NoRot*) were rotated back into alignment with the scanner axes (see black arrow in Figure 2). Unless stated otherwise, all processing and analysis operations were carried out using MATLAB (MathWorks Inc., Natick, MA, USA).

### Numerical Phantom Investigations

Multi-echo (*TE* = 4,12, 20, 28 ms) magnitude and phase images, from a numerical phantom^21^, with (originally) no phase wraps or background fields present, were used to independently investigate the effect of the three tilt correction methods (described above), and no correction, on each step in the QSM pipeline.

We carried out these investigations with two image volumes: one unpadded with the original matrix size l64×205 ×205 and a second volume padded to 357 × 357 × 357. The padded matrix size was chosen as the long diagonal of the initial volume (padded to a cube: 205 × 205 × 205) and rounded up to the nearest odd integer. This was to ensure that none of the original frequency coefficients of the unit dipole field were cut off due to rotations about any of the three axes. An odd matrix size meant that there was a true centre of rotation correctly located within a single central voxel.

#### Numerical Phantom: Susceptibility Calculation

Local field maps, Δ*B_int_*(***r***), were calculated from a non-linear fit^22^ over all echo times (for the most accurate field estimates^23^) of the complex data set created by combining the magnitude and background field-free phase images. These local field maps, obtained from the supplied raw numerical phantom data, were free of any (synthetic) background fields or phase wraps, and therefore allowed investigation of the effect of oblique acquisition on *χ* calculation alone. To simulate oblique acquisition, local field maps were rotated between ±45° in 5° increments. All tilt correction methods described (and no correction) were compared for three *χ* calculation methods chosen to cover the two main approaches: direct non-iterative solutions (in *k*-space) and iterative solutions (in image-space).

The first method tested was direct, thresholded *k*-space division (TKD)^24,25^ (from open-source software) where a modified dipole kernel was generated in *k*-space with values below a threshold, δ = 2/3, replaced by the signed threshold value:

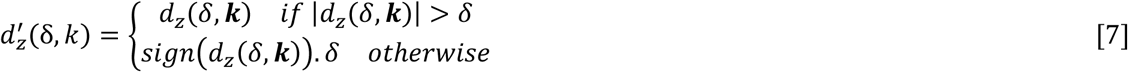

The dipole was originally defined according to *DipK* and *DipIm* and then always thresholded in *k*-space. Susceptibility underestimation was corrected by multiplication with a correction factor, *c_χ_*(*δ*), calculated according to^26^.

The second and third *χ* calculation methods aim to iteratively solve for *χ* through the minimisation of

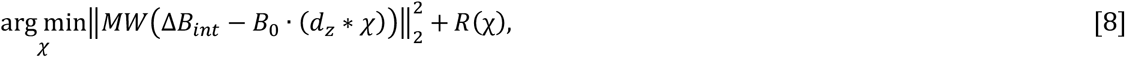

where *M* is a binary mask, *W* is a weighting term and *R* (χ) is the data regularisation term that reflects some prior information about χ. Iterative Tikhonov regularisation^25,27^ (open-source) was chosen as it has performed well in a variety of QSM applications including outside the brain^28–30^. It was applied with 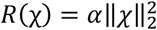, a regularisation parameter α = 0.003 (chosen through an L-Curve analysis^31^), and W reflecting the spatially varying noise, and was also corrected for *χ* underestimation^26^. Weighted linear total variation regularisation (from the FANSI toolbox^18,32^) with *R*(χ) = *α*|∇_*χ*_|_1_, *α* = 6.31 × 10^-5^ (chosen through an L-Curve analysis) and W the magnitude of the complex data^32^ was also tested. This method was chosen as total variation based iterative approaches were shown to produce the most accurate susceptibility maps in the 2019 QSM challenge 2.0^33^.

Mean χ values were calculated in five deep gray matter regions of interest (ROIs): the caudate nucleus, globus pallidus, putamen, thalamus and red nucleus. All susceptibility maps were compared using the root mean square error (RMSE) and QSM-tuned structural similarity index (XSIM)^34^ metrics relative to the supplied ground truth susceptibility map at 0°.

#### Numerical Phantom: Background Field Removal

For the background field removal step, local field maps from the numerical phantom required the addition of synthetic background fields, which were then removed following the three different tilt-correction methods (and no correction). After background field removal, the susceptibility maps were calculated from the resulting field maps using the *χ* calculation method found to be optimal in the above described assessment.

To investigate the effect of tilt correction schemes on the background field removal step, synthetic background fields, Δ*B_ext_*(***r***) (Figure 3, bottom left), were added to the local field maps used in the *Methods: Numerical Phantom: Susceptibility Calculation* section. The background fields were calculated using the forward model, i.e. through a convolution, formulated as a multiplication in Fourier space, between the unit magnetic dipole field and a head-shaped susceptibility map^16,17^:

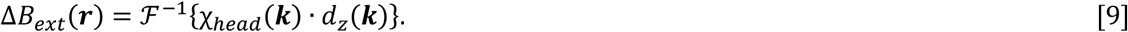

**Figure 3:**
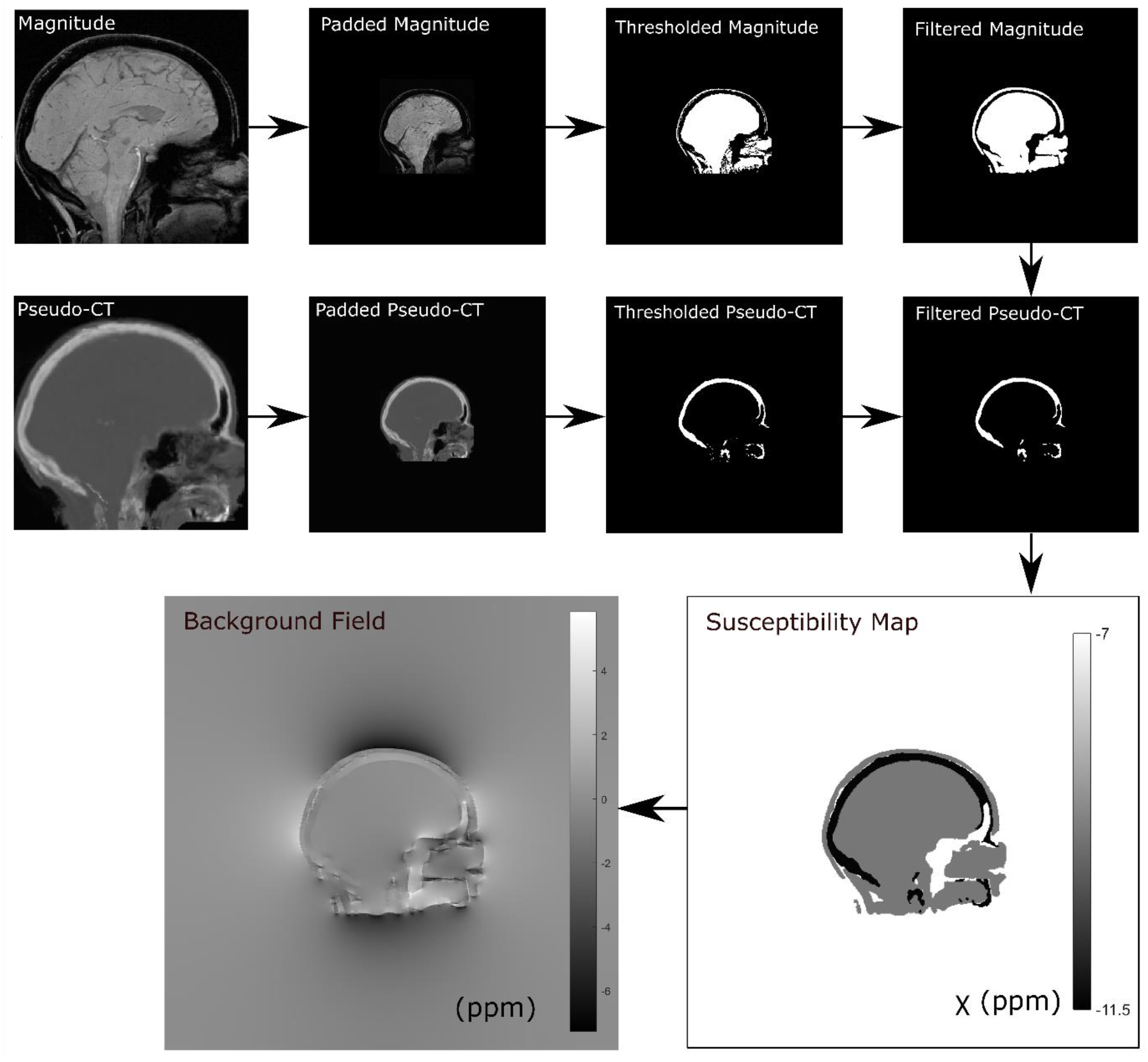
Method for calculating the synthetic background field from a head-shaped susceptibility map obtained by thresholding the numerical phantom magnitude image and a pseudo-CT image to delineate soft tissue and bone, respectively. The thresholded magnitude and pseudo-CT images were filtered for smoothness using a 3 × 3 × 3 box-filter.

*χ_head_* is the head-shaped susceptibility map, with soft tissue (−9.4ppm) and bone (−11.4ppm)^7,35^ regions obtained by thresholding the magnitude (sum of squares over all echoes) and a pseudo-CT^36–38^ image, respectively (Figure 3). The magnitude and pseudo-CT images were padded from their original matrix size of 164 × 205 × 205 to 512 × 512 × 512 to ensure edge effects from the periodic Fourier transform were minimised around the volume of interest. These synthetic background fields were then cropped back to their original matrix size and added to the local field maps obtained previously simulating a total field map, Δ*B*(***r***) = Δ*B_int_*(***r***) + Δ*B_ext_*(***r***). To simulate oblique acquisition, total field maps were rotated between ±45° in 5° increments.

To remove the synthetic background fields, Δ*B_ext_*(***r***), from the tilted total field maps three different state-of-the-art background field removal methods^39^ were used, based on their widespread use, robustness and accuracy^39,40^. Projection onto dipole fields (PDF)^41^, from the MEDI Toolbox^14^, was used following tilt correction with all three correction schemes and no correction because PDF is orientation dependent, i.e. it uses the dipole field *d_z_*(***k***) (Equation 5). Laplacian boundary value (LBV)^42^, from the MEDI Toolbox^14^, and variable-kernel sophisticated harmonic artifact reduction for phase data (V-SHARP)^43^, from STI Suite^19^, were tested with *RotPrior* and *NoRot* only, as LBV and V-SHARP are orientation independent methods, i.e. they do not use the dipole field. Following rotation back into the reference frame (equivalent to *RotPrior* for the susceptibility calculation step), susceptibility maps were calculated from all local field maps using iterative Tikhonov regularisation (regularisation parameter *α* = 0.003) as this was found to be optimal. Susceptibility maps were compared using RMSE and XSIM^34^ relative to the ground truth susceptibility map at 0°.

#### Numerical Phantom: Phase Unwrapping

To investigate the effect of tilt correction on phase unwrapping, the synthetic background fields added in the previous section (*Methods: Numerical Phantom: Background Field Removal*) induced phase wraps when the phase was constrained to the [–π, π) interval, which were then unwrapped. Susceptibility maps were then calculated from these unwrapped field maps using background field removal and susceptibility calculation algorithms found to be optimal in the experiments described in the previous sections (*Methods: Numerical Phantom: Susceptibility Calculation* and *Background Field Removal*).

To investigate the effect of tilt correction on phase unwrapping, phase wraps were introduced into the wrap-free numerical phantom images via the additional synthetic background field described above. From each total field map at each angle, Δ*B*(***r***) = Δ*B_int_*(*r*) + Δ*B_ext_*(***r***), multi-echo unwrapped phase images were simulated via scaling the tilted total field maps at each echo time according to *ϕ*(***r**, TE*)=*γ*·*TE* · Δ*B*(***r***).

At every tilt angle, a complex data set (*S*) was made from the multi-echo magnitude images (*M*) and simulated phase images (*ϕ*) using 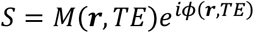, which constrained the phase to the range [—π, π), resulting in phase wraps. A wrapped total field map was calculated via a non-linear fit over all echo times^22^, which then underwent phase unwrapping using the commonly used Laplacian^14,44^, SEGUE^45^ (https://xip.uclb.com/product/SEGUE) and ROMEO^46^ (https://github.com/korbinian90/ROMEO/releases) techniques with the *NoRot* and *RotPrior* tilt correction methods. After rotating all the unwrapped images back into the reference frame (equivalent to *RotPrior* for the background field removal step and susceptibility calculation step), susceptibility maps were then calculated with PDF^41^ background field removal and susceptibility calculation using iterative Tikhonov regularisation (regularisation parameter *α* = 0.003) as we found these to provide optimal results.

### Investigations In Vivo

#### In Vivo: MRI Acquisition

3D gradient-echo brain images of five healthy volunteers were acquired on a 3T Siemens Prisma-Fit MR system (National Hospital for Neurology and Neurosurgery, London, UK) using a 64-channel head coil across a range of image volume orientations. Note that the volunteers did not tilt their head but remained in the same position throughout the experiment. The image volume was tilted about the x-axis, as this is the most common in clinical practice, from −45° to +45° in 10° increments, with the reference image at 0° representing a non-oblique acquisition (subject 5 angles were between ±15°). Each image volume was acquired in 3 min 23 s with TR = 30ms; TEs = 4.92, 9.84, 14.76, 19.68, 24.60 ms; 1.23 mm isotropic voxels; FOV = *256 × 192 × 216.6* mm; Matrix Size = *208* × 156 × 176; bandwidth = 280 Hz/pixel; flip angle 15°; 6/8 partial Fourier along PE1 and PE2; on the scanner ASPIRE coil combination^47^; monopolar readout; and GRAPPA_PE1_ acceleration = 3 (FE direction: A>>P, PE1 direction: R>>L, PE2 direction F>>H).

#### In Vivo: Phase Unwrapping

For all angles/volumes, a total field map and a noise map were obtained using a non-linear fit of the complex data^22^ from the MEDI toolbox^14^. A brain mask was created using the brain extraction tool (BET)^48^ with default settings applied to the final echo magnitude image of the reference 0° volume for a conservative brain mask estimate. This brain mask was then registered to all oblique acquired volumes to maintain consistency. As with the numerical phantom, both the *RotPrior* and *NoRot* correction schemes were applied. Residual phase wraps were then removed using Laplacian unwrapping^44^, SEGUE^45^ and ROMEO^46^. To investigate the effect of the correction schemes on this step in the pipeline, unwrapped total field maps were rotated back into the reference frame (equivalent to *RotPrior* for the background field removal step and susceptibility calculation step) and susceptibility maps were created using PDF background field removal and susceptibility calculation with iterative Tikhonov regularisation (α *= 0.017* chosen through an L-Curve analysis).

As in the numerical phantom, and also due to very slight unavoidable changes in subject position between scans, the unwrapped field maps and susceptibility maps were registered into the reference image space to facilitate comparisons of results in vivo. To carry out this registration, the magnitude image (added in quadrature over all echoes) for each angle was rigidly registered to the 0° magnitude using NiftyReg^49^ resulting in a transformation matrix per angle/volume, which was applied to bring all angles/volumes into the same common reference space.

#### In Vivo: Background Field Removal

The ROMEO unwrapped field maps described above (in *Methods: <u>In Vivo: Phase Unwrapping</u>*) for volumes at all angles, and prior to any registrations or rotations, were used to investigate the effect of oblique acquisition on background field removal. ROMEO was chosen as it has been shown to outperform^46^ PRELUDE^50^ and BEST-PATH^51^. As for the numerical phantom, for each field map, at each angle, background fields were removed using PDF^41^ with all tilt correction schemes and no correction, and using LBV^42^ and V-SHARP^43^ with only *RotPrior* and *NoRot*. For all three background field removal methods, the brain mask was eroded by 4 outer voxels^52^. *RotPrior* was performed twice: with mask erosion either before or after the rotation to compare the effects of interpolation, particularly along the boundaries of the field map on PDF and V-SHARP, as it is known that boundary effects arise in these background field removal methods^39^.

For comparison purposes, after rotation and registration of the local field maps back into the reference frame (equivalent to *RotPrior* for the susceptibility calculation step), susceptibility maps were calculated from the local field maps using iterative Tikhonov regularisation (α = 0.017, chosen with an L-Curve). Local field maps and susceptibility maps were compared with RMSE and XSIM metrics (XSIM only for the susceptibility maps) averaged across all subjects relative to the 0° reference image.

#### In Vivo: Susceptibility Calculation

To investigate the effect of oblique acquisition on the χ calculation step in the pipeline, we used the local field maps following ROMEO unwrapping and LBV background field removal (described in the previous section *Methods: In Vivo: Background Field Removal*), prior to any registrations or rotations. LBV was chosen as it is orientation independent, thereby allowing our analysis to focus on the effect of oblique acquisition and the different correction schemes on the χ calculation step alone. The three tilt correction schemes (and no correction) were compared using the same three χ calculation methods as for the numerical phantom: TKD, iterative Tikhonov regularisation with a regularisation parameter α = 0.017 from an L-curve analysis, and weighted linear TV with a regularisation parameter α = 6.31 × 10^-5^ found also from an L-curve.

The resulting susceptibility maps were transformed into the reference space as described in the earlier section *Methods: In Vivo: Phase Unwrapping*. The same ROIs as in the numerical phantom were investigated and were obtained by registering the EVE^53^ magnitude image with the reference magnitude image (at the first echo time) and applying the resulting transformation to the EVE ROIs. Mean *χ* values were calculated in these ROIs for all tilt angles and all correction schemes. RMSE and XSIM measures averaged across all subjects were also used to compare the susceptibility maps.

## Results

### Numerical Phantom

All numerical phantom results shown here are for rotations of image volumes about the *x*-axis with unpadded matrices. Note that acquisitions tilted about the *y*-axis and *y*=*x* axis as well as images with padded matrices all gave similar results (see Supporting Figures S1–3). We chose to display these results as the padded matrix size leads to increased computation time, which is not recommended in a practical setting, and the *x*-axis is the most common axis of rotation for oblique acquisition.

When wrapped phase images are rotated prior to phase unwrapping with the correction scheme *RotPrior*, artefacts arise for both Laplacian and SEGUE unwrapping (Figure 4). SEGUE appears to fail with *RotPrior* as it incorrectly identified phase wraps and thereby removed a portion of the mask (Figure 4c).

**Figure 4:**
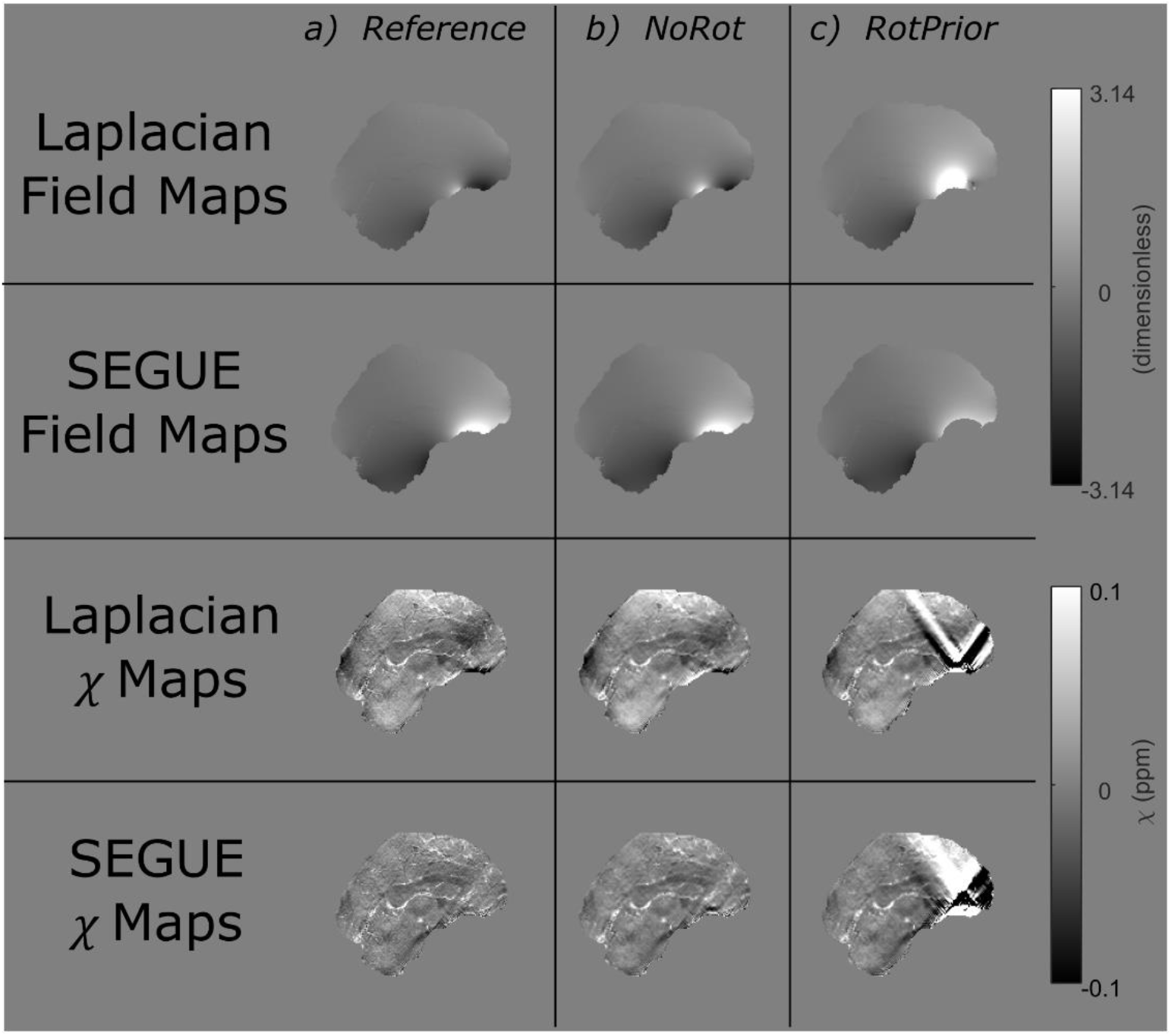
The effect of tilt correction before phase unwrapping in the numerical phantom. Phase unwrapped field maps and the resulting susceptibility maps at 15° for the *NoRot* (column b) and *RotPrior* (column c) tilt correction methods relative to the reference (column a). Rotation of the wrapped field maps prior to phase unwrapping with both Laplacian and SEGUE techniques results in errors along phase wraps and incorrect unwrapping, leading to prominent artefacts in the final susceptibility maps.

When using PDF for background field removal, *RotPrior* is the most accurate method, and the largest errors arise from *DipIm* and *NoRot* (Figure 5). Striping artefacts are present in the local field map from the *DipK* method. LBV and V-SHARP are shown to be largely unaffected by oblique acquisition.

Figure 6 summarises the mean susceptibility in the Caudate Nucleus and Thalamus, alongside XSIM measurements across all angles for all three *χ* calculation methods (RMSE results are shown in Supporting Figure S4, and are similar). TKD and iterative Tikhonov methods are most accurate with *RotPrior*, and least accurate when the dipole is misaligned to the main magnetic field (*NoRot*). Weighted linear TV is relatively robust to oblique acquisition with *RotPrior* and *DipK* performing similarly. However, *DipK* shows more variability in *χ* at the ROI level than *RotPrior*. Weighted linear TV with *DipIm* fails at non-zero angles and *NoRot* results in the largest errors. Example susceptibility maps are shown in Figure 7, highlighting the widespread *χ* errors that arise when the magnetic dipole is defined incorrectly (*NoRot*).

**Figure 5:**
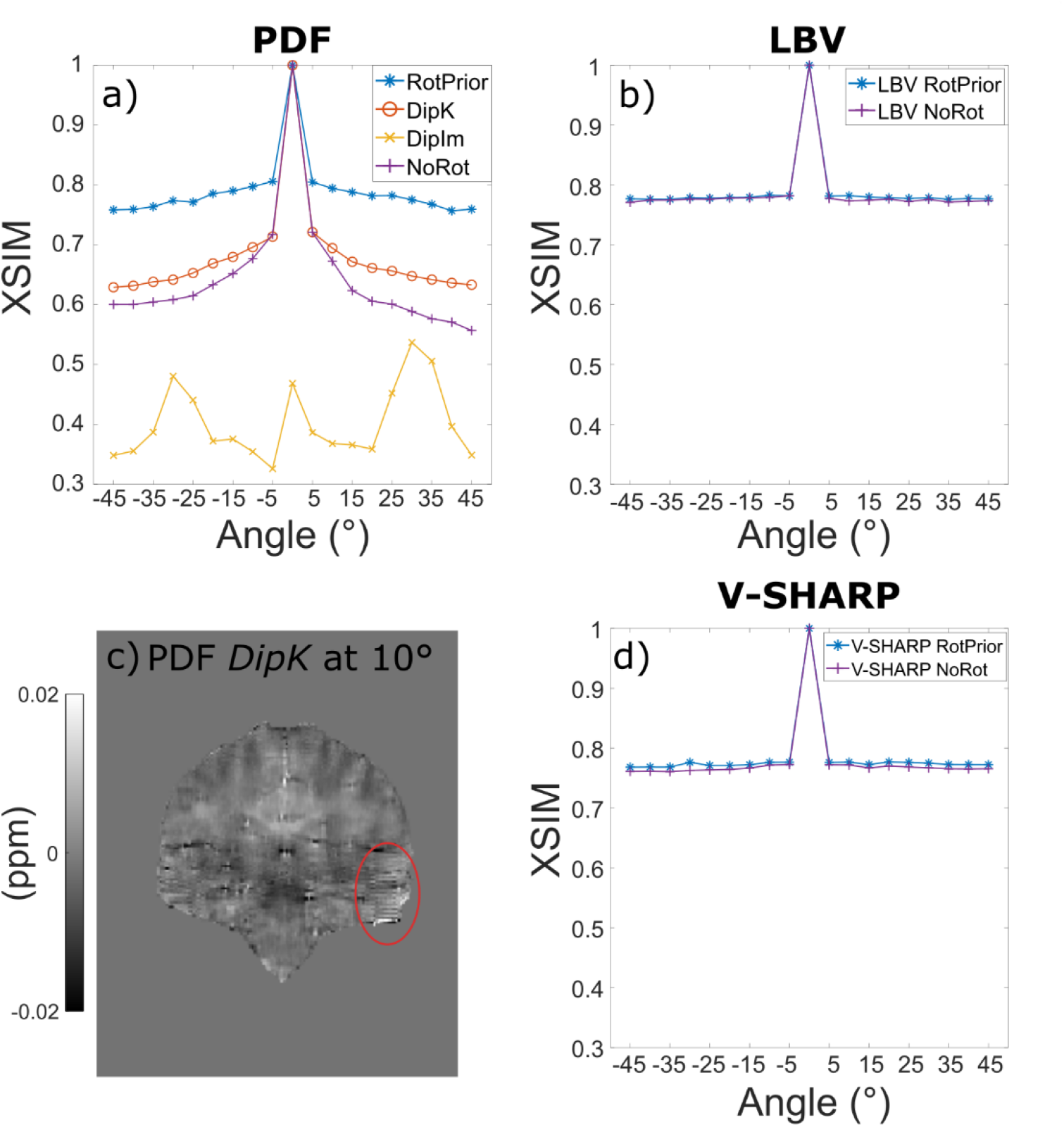
The effect of different tilt correction schemes on QSM with three background field removal methods in a numerical phantom. Susceptibility maps for XSIM comparisons were calculated with iterative Tikhonov regularisation. For PDF (a) the XSIM metric shows that *RotPrior* gives the most accurate susceptibilities, with *DipIm* performing the worst. When using PDF with *DipK*, striping artefacts (c, red ellipse) arise in the local field maps for tilted acquisitions. LBV and V-SHARP (b, d) are shown to be largely unaffected by oblique acquisition with differences arising primarily from rotation interpolations.

**Figure 6:**
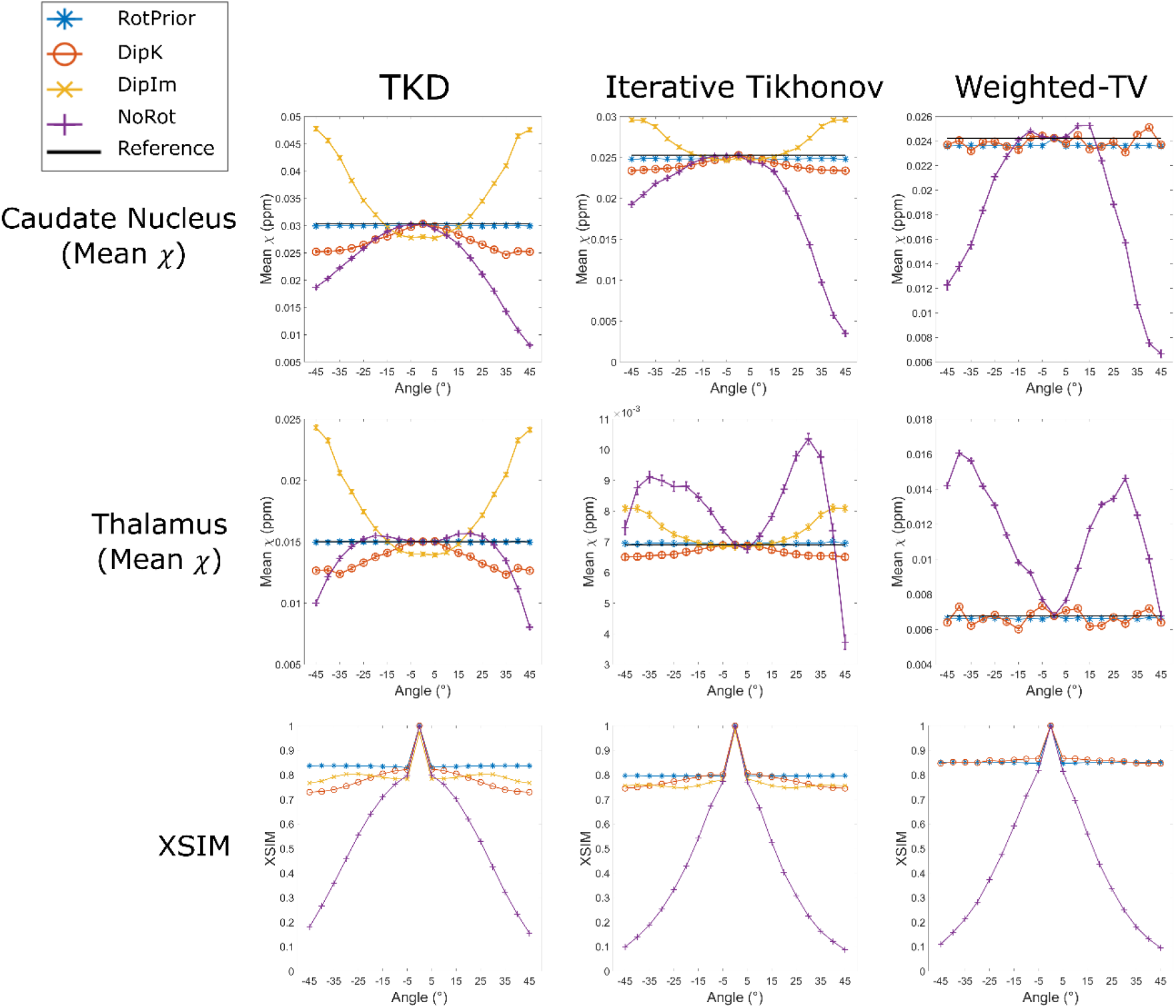
Mean susceptibilities in the caudate and thalamus (top rows), and XSIM (bottom row) across all tilt angles for all tilt correction schemes and all three *χ* calculation methods in the numerical phantom. RMSE measurements shown in Supporting Figure S4 agree with XSIM findings. *NoRot* performs worst across all angles. *RotPrior* is the most accurate tilt correction scheme. For weighted linear TV, *DipK* and *RotPrior* have similar XSIM values but the mean thalamus *χ* varies more over angles with *DipK*. Note that *DipIm* is not shown for weighted linear TV as this method fails.

**Figure 7:**
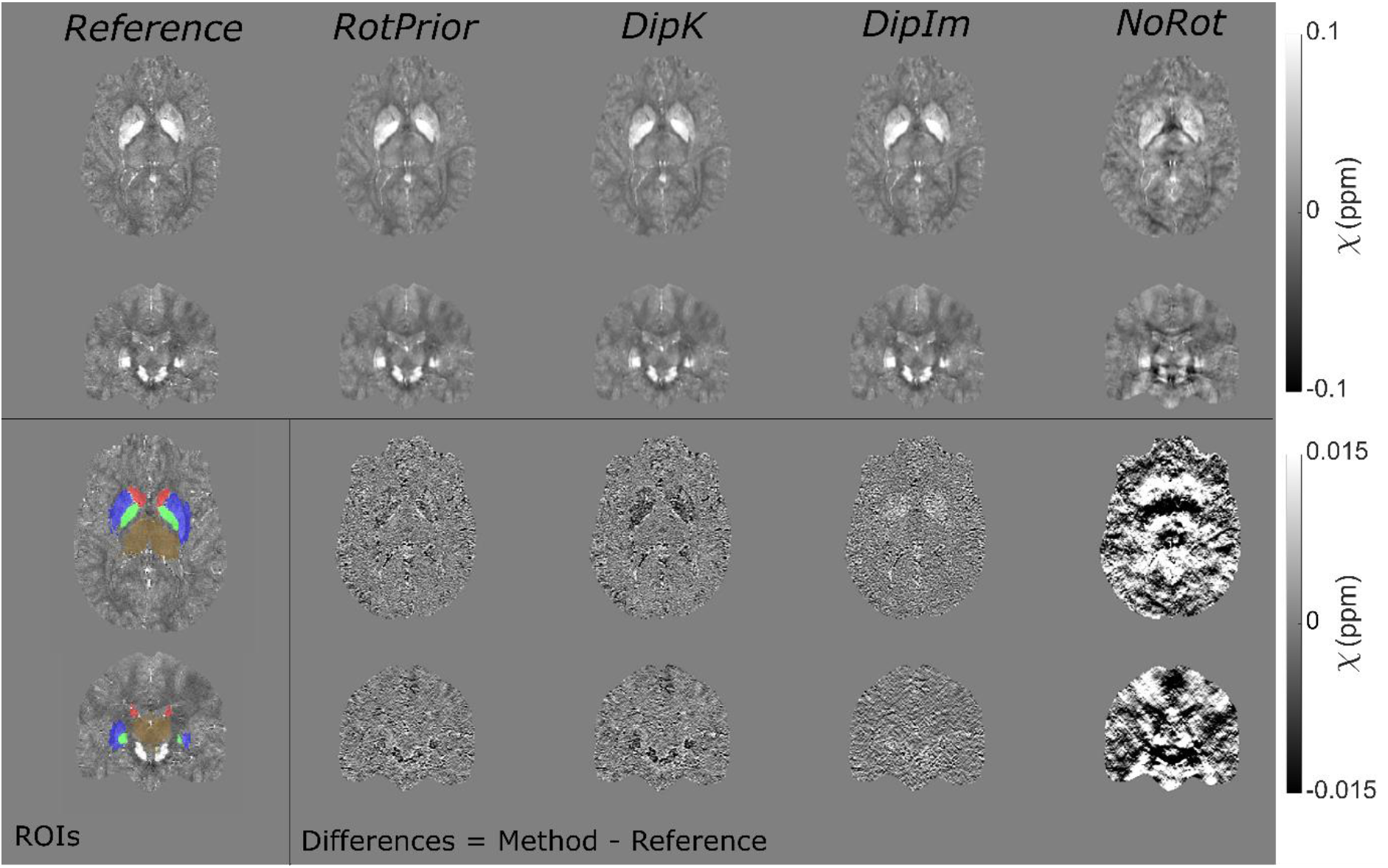
*χ* maps and difference images illustrating the effects of all tilt correction schemes in the numerical phantom. An axial and a coronal slice are shown for a volume tilted at 25° and a reference 0° volume with all *χ* maps calculated using the iterative Tikhonov method. The ROIs analysed are also shown (bottom left). *RotPrior* performs the best while *NoRot* results in substantial *χ* errors across the whole brain. The results from TKD and weighted linear TV (not shown) are very similar.

### In Vivo

In vivo, Laplacian and SEGUE phase unwrapping with *RotPrior* have the same image artefacts as in the numerical phantom (not shown here) compared with *NoRot*, with incorrect identification of phase wraps when wrapped field maps are rotated prior to phase unwrapping.

Figure 8 shows that PDF background field removal is most accurate with *RotPrior* and least accurate with *DipIm* followed by *NoRot*, confirming the results obtained in the numerical phantom (Figure 5). The average XSIM differences between tilt correction schemes in vivo (Figures 8 and 9) are smaller than in the numerical phantom (Figures 5 and 6), most likely due to issues inherent to in-vivo acquisition including motion and much greater noise. Striping artefacts are present in the *DipK* method for PDF in the local field maps prior to re-orientation for comparison purposes (Fig. 8c). Rotation interpolation obscures these artefacts in the in-vivo images. LBV and V-SHARP are shown to be largely orientation independent in the in vivo case as expected.

**Figure 8:**
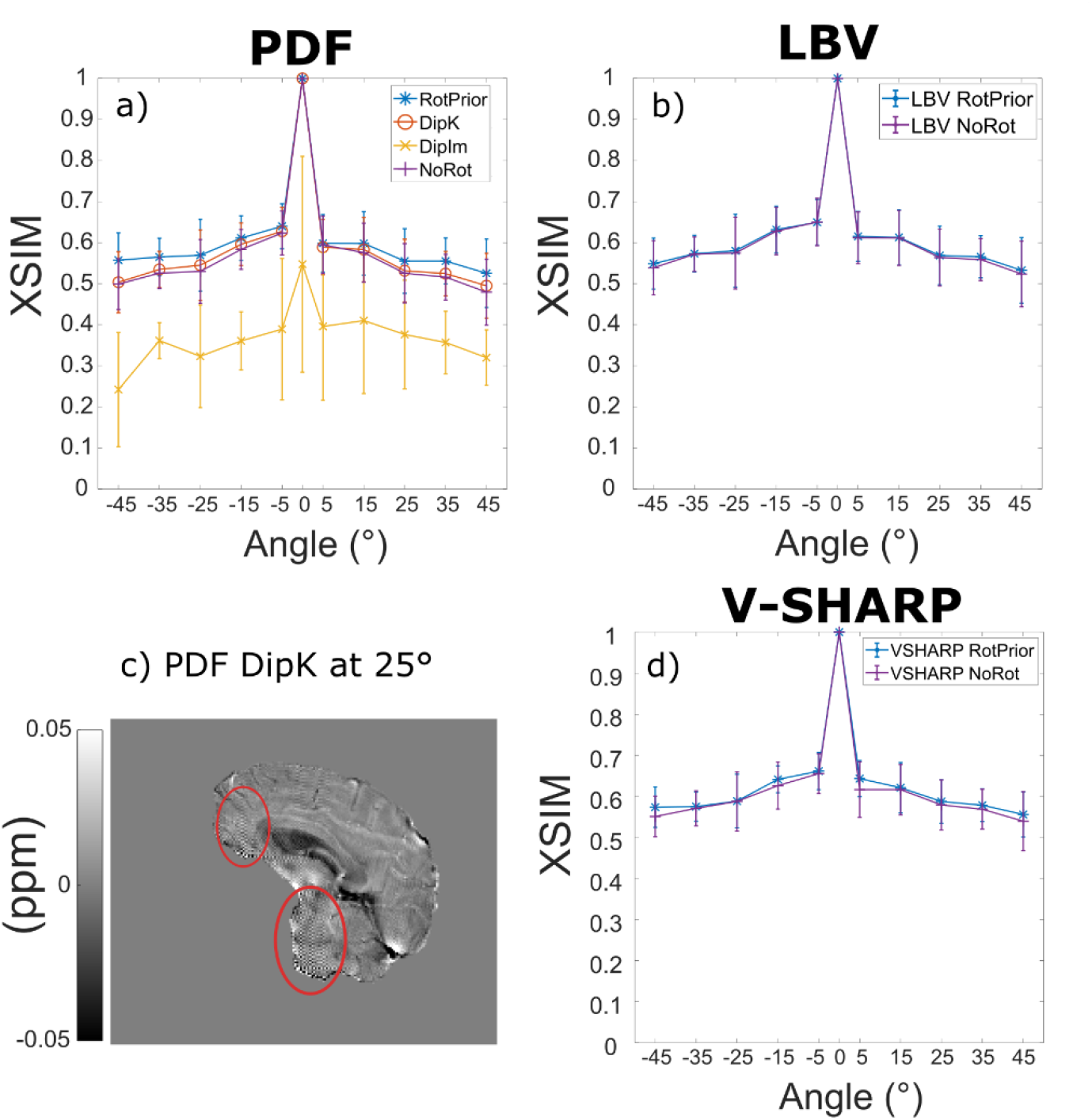
The effect of different tilt correction schemes on background field removal in vivo. Average XSIM measurements across all subjects were used to compare the *χ* maps calculated with iterative Tikhonov regularisation after background field removal to the non-oblique (0°) reference map. PDF (a) has the highest XSIM with *RotPrior* and the lowest XSIM with *DipIm*, followed by *NoRot*, confirming the results in the numerical phantom (Figure 5). Striping artefacts are found in local field maps when using *DipK* and PDF (c, red ellipses) but are obscured after rotation and registration back into the reference 0° space due to interpolation. LBV (b) and V-SHARP (d) are shown to be unaffected by oblique acquisition in vivo as well as in the numerical phantom (Figure 5d). Error bars represent the standard deviation of the mean XSIM across subjects.

**Figure 9:**
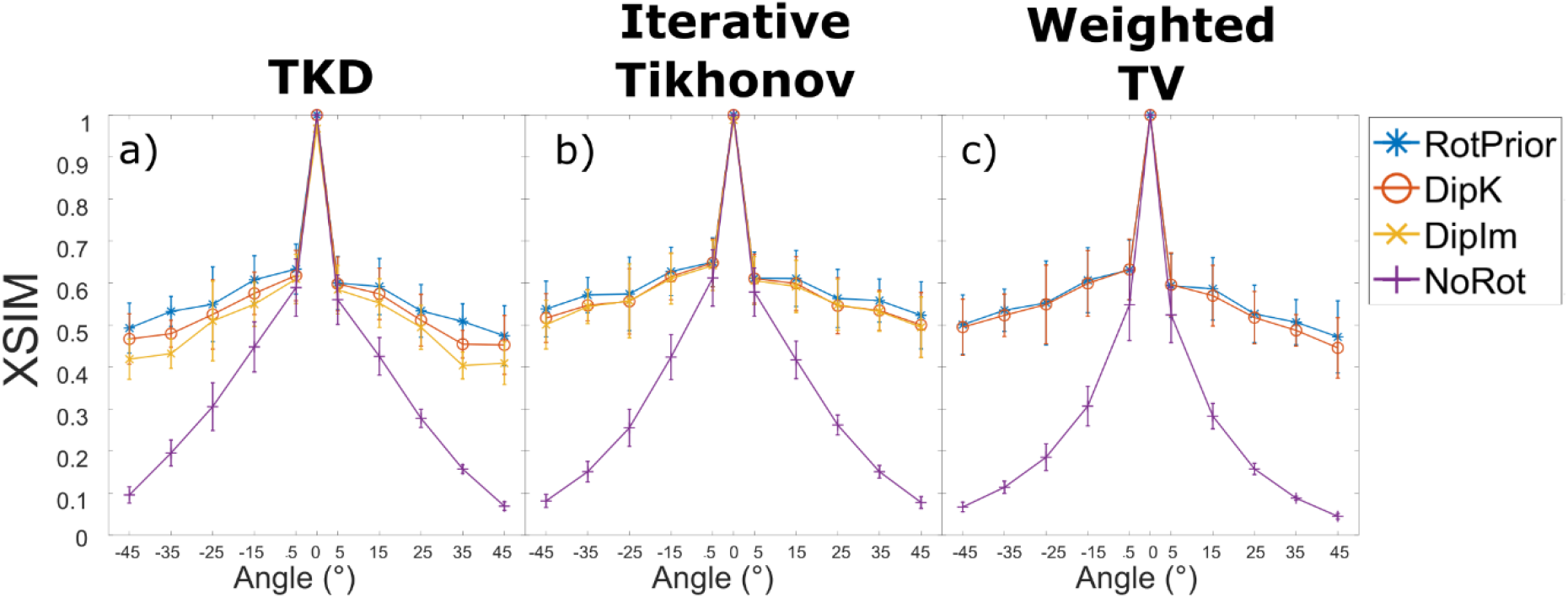
Average XSIM plots over all angles for all tilt correction schemes and all three *χ* calculation methods across all subjects in vivo. RMSE measurements in Supporting Figure S4 agree with these XSIM findings. These results are similar to those in the numerical phantom (Figure 3) with *RotPrior* consistently reporting higher XSIM measures than other methods and *NoRot* performing worst across all methods. At non-zero tilt angles, XSIM have a respectively high/low baseline level arising from rotation and registration interpolations. *DipIm* fails for weighted linear TV and is, therefore, omitted from the plots in the last column. Error bars represent the standard deviation of the mean XSIM across subjects.

When *RotPrior* was performed with mask erosion before rotating the total field map, artefacts arose along the boundaries of the PDF local field map (See Supporting Figure S5). PDF performs more robustly if mask erosion is carried out after rotation, whereas V-SHARP appears to perform equally well in both scenarios.

Figure 9 shows the effect of all tilt correction schemes on susceptibility calculation in vivo and confirms that *NoRot* results in the largest susceptibility errors and that *RotPrior* is consistently the most robust tilt correction method compared to other methods. Both *RotPrior* and *DipK* perform better than *DipIm*, in agreement with results in the numerical phantom (Figure 6). Difference images (Figure 10) also confirm those obtained in the numerical phantom (Figure 7). Subtle effects found in several of the numerical phantom ROIs (Figure 6) were not apparent in vivo.

**Figure 10:**
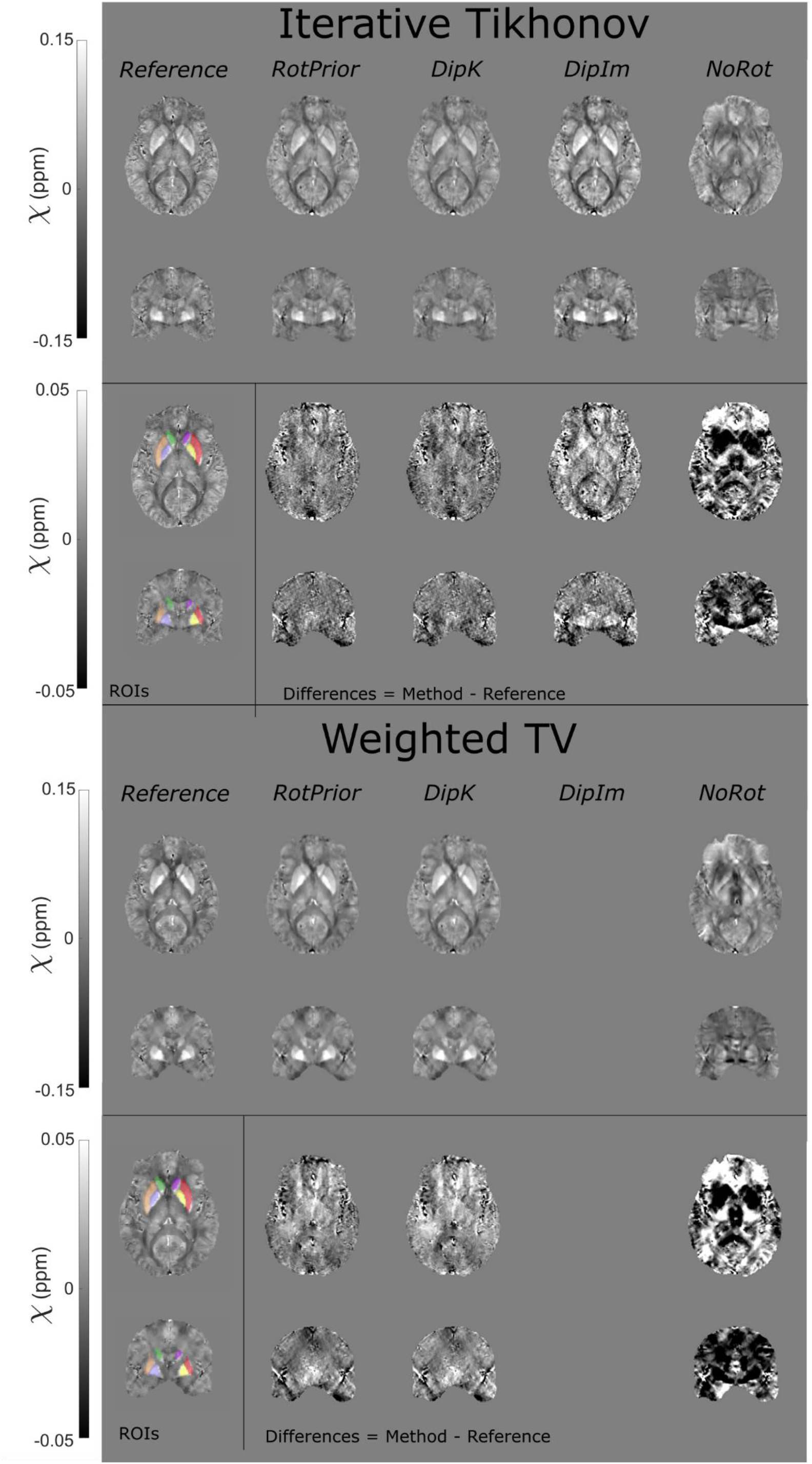
χ maps and difference images illustrating the effects of all tilt correction schemes on susceptibility calculation in vivo. An axial and a coronal slice are shown for a volume tilted at 45° and a reference (0°) volume with all χ maps calculated using the iterative Tikhonov method (top) and weighted linear TV (bottom). Weighted linear TV with *DipIm* fails at non-zero angles and is therefore omitted from the figure. *NoRot* leads to the largest differences and image artefacts throughout the brain for iterative Tikhonov and weighted linear TV methods. The EVE ROIs used are shown (bottom left). Results from TKD (Supporting Figure S6) are very similar.

## Discussion

We have shown that oblique acquisition must be accounted for in the QSM pipeline to ensure accurate susceptibility estimates throughout the brain. For all background field removal and susceptibility calculation methods tested, if the magnetic dipole kernel is left misaligned to the 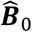 direction (*NoRot*), which can arise from user error in popular QSM toolboxes, then significant susceptibility errors result.

Through the analysis of the effect of tilted acquisition on a numerical phantom and five healthy volunteers in vivo, we have shown that any rotations that are applied to a wrapped field map prior to phase unwrapping will result in incorrect unwrapping, using Laplacian and SEGUE unwrapping techniques, and subsequent artefacts in the resulting QSMs. Results indicate that, for PDF background field removal, rotating the image into the scanner frame and using a *k*-space dipole defined in the scanner frame (*RotPrior* correction method) provides the most accurate susceptibility maps. If no image rotations are desired, due to unwanted interpolation effects, LBV or V-SHARP are recommended as they are largely unaffected by oblique acquisition. Both TKD and iterative Tikhonov susceptibility calculation methods provide the most accurate results when local field maps are rotated into alignment with the scanner axes and a *k*-space dipole, defined in the scanner frame, is used (*RotPrior*). The same conclusion holds for weighted linear TV, but susceptibility calculation can be carried out in the oblique image frame without any rotations provided the correct ***B***_0_ direction is used in defining the *k*-space dipole (*DipK* correction method). We therefore recommend rotating the total field map into alignment with the scanner frame after phase unwrapping but prior to background field removal.

Both the numerical phantom and in vivo results indicate that when wrapped phase images are rotated prior to phase unwrapping (with the correction scheme *RotPrior*), artefacts arise for both Laplacian and SEGUE unwrapping methods. This is probably due to interpolation errors along phase wraps (Figure 4). Therefore, any phase unwrapping must be carried out in images left in the same orientation as acquired, with rotations only being applied afterwards to avoid artefacts.

When using PDF for background field removal, numerical phantom and in vivo results show that *RotPrior* consistently provides the most accurate susceptibility maps while *NoRot* performs the worst (Figures 5 and 8). Striping artefacts arise in local field maps in both the numerical phantom and in vivo when using PDF with the *DipK* method. First identified by Dixon, E.^7^, these striping artefacts are present due to the violations in circular continuity when defining the tilted dipole in *k*-space and using the inverse discrete Fourier transform to transform the susceptibility maps into image-space (which enforces periodicity, see Supporting Figure S7). Striping artefacts arise from regions of high susceptibility changes, such as on the brain boundaries (Figures 5c and 8c). *DipIm* also resulted in poor background field removal, most likely due to fitting to the incorrect twisted or sheared unit dipole field (bottom row of Figure 1). To avoid artefacts and robustly achieve background field removal with PDF, total field maps must be rotated into alignment with the scanner frame prior to PDF background field removal as it is then possible to use the non-oblique dipole, which doesn’t violate circular continuity (Supporting Figure S8). We showed that LBV and V-SHARP were mostly unaffected by oblique acquisition, with the differences between zero and non-zero tilt angles arising solely from rotation interpolation effects.

Given that *RotPrior* is the most accurate method for PDF, the typically necessary mask erosion must be carried out after rotation into the reference space. Artefacts that arise along the boundaries of the local field map if erosion is carried out prior to rotation (Supporting Figure S5a and S5b) probably arise from distortion of the dipolar background fields due to interpolation at the edges. In contrast, V-SHARP does not show substantial differences with mask erosion before *v*. after rotation which suggests that the interpolation may not substantially affect the harmonic nature of the background fields on which this method relies.

We found TKD and iterative Tikhonov regularisation to be affected by oblique image orientation and most accurate with *RotPrior*. We showed weighted linear TV to be relatively robust to oblique acquisition, however *RotPrior* is still maintained to be the most consistently robust method (Figure 9). For all susceptibility calculation methods tested, a unit dipole field misaligned to the main magnetic field (*NoRot*) leads to artefacts and substantial errors in susceptibility maps. The subtle differences between correction methods found in the numerical phantom ROIs (Figure 6) were not apparent in vivo probably due to noise, motion and the expected variability in susceptibility maps over repeated acquisitions^27,54^.

At non-zero tilt angles, XSIM (and RMSE in Supporting Figure S4) values have a respectively high (or low) baseline level arising from rotation (no matter how small the angle) and registration interpolations, and imperfections inherent to in vivo acquisition. Additional discrepancies in similarity measures between rotated and unrotated QSM may also have occurred due to slight differences in repeated acquisitions as evidenced by the slightly larger (0° to 5°) XSIM discrepancy in vivo than in the numerical phantom (Figures 8 and 9 compared with Figures 5 and 6). The effect of these discrepancies has been minimised by averaging across all five healthy volunteers.

Our results also indicate that at larger tilt angles in the numerical phantom, *DipK* is less accurate than *RotPrior*, therefore, for certain imaging applications including cardiac imaging and pelvic imaging where large tilt angles of up to 45° are often required, tilt correction is likely to be essential for accurate susceptibility mapping.

Therefore, we recommend accounting for oblique acquisition by using the *RotPrior* tilt-correction method before background field removal since this method gave the most accurate susceptibility maps in both the numerical phantom and in vivo. If desired, the susceptibility map can be rotated back into the original orientation after susceptibility calculation to facilitate comparison with other (processed) images. In the future, it would be interesting to carry out a similar investigation of tilt correction methods for total field inversion (TFI)^55,56^, in which background field removal and susceptibility calculation are combined into a single step. Due to the reliance of TFI on the correct definition of the magnetic dipole, we would expect to see similar results, with *RotPrior* being more robust to oblique acquisition. However, this would need to be confirmed with further work. It is possible to build an alternative rotation-free pipeline of methods relatively unaffected by oblique acquisition (such as LBV and weighted linear TV) but those methods must be checked to ensure true independence of image orientation. However, such an approach limits the choice of methods for the steps in the QSM pipeline, which could lead to suboptimal susceptibility maps. For example, LBV’s highly specific boundary approximations can be easily violated making it easier to simply rotate the field maps in some cases. These aspects must be considered carefully when designing a QSM pipeline.

## Conclusions

Oblique acquisition must be accounted for in the QSM pipeline to avoid artefacts and erroneous susceptibility estimates. We recommend rotating the total field map into alignment with the scanner frame after phase unwrapping but before background field removal (and then rotating the final susceptibility map back into the original orientation). Alternatively, a QSM pipeline relatively robust to oblique acquisition can be built from a more limited number of image-orientation-independent methods (e.g. LBV or V-SHARP for background field removal and weighted linear TV for susceptibility calculation). However, care must be taken in weighing up the minimal effects of image interpolation (from tilt correction rotations) versus choosing from a smaller range of methods that are orientation independent and may not be as robust oblique acquisition, as they may not be as accurate nor optimal for a given data set. It would also be vital to ensure a chosen method is independent of slice orientation, which may require further investigation. Our recommended correction scheme ensures that all methods developed for each stage of the QSM pipeline can be used and optimised.

We provide an open-source MATLAB function that can be easily incorporated into existing MATLAB-based QSM pipelines and used both to align oblique 3D volumes with the scanner frame and to rotate image volumes (e.g., susceptibility maps) back into the original oblique image orientation. This function is freely available at: https://github.com/o-snow/QSM_TiltCorrection.git. Our results allow accurate susceptibility maps to be obtained from oblique image acquisitions. This is an important step in translating QSM into clinical practice.

## Acknowledgments

Oliver Kiersnowski’s work was supported by the EPSRC-funded UCL Centre for Doctoral Training in Intelligent, Integrated Imaging in Healthcare (i4health) (EP/S021930/1). John Thornton received support from the National Institute for Health Research University College London Hospitals Biomedical Research Centre. Karin Shmueli and Anita Karsa were supported by European Research Council Consolidator Grant DiSCo MRI SFN 770939. We would like to thank Carlos Milovic for his help with the FANSI toolbox, and the healthy volunteers for their time.

## Supporting Information Figures

**Supporting Figure S1:**
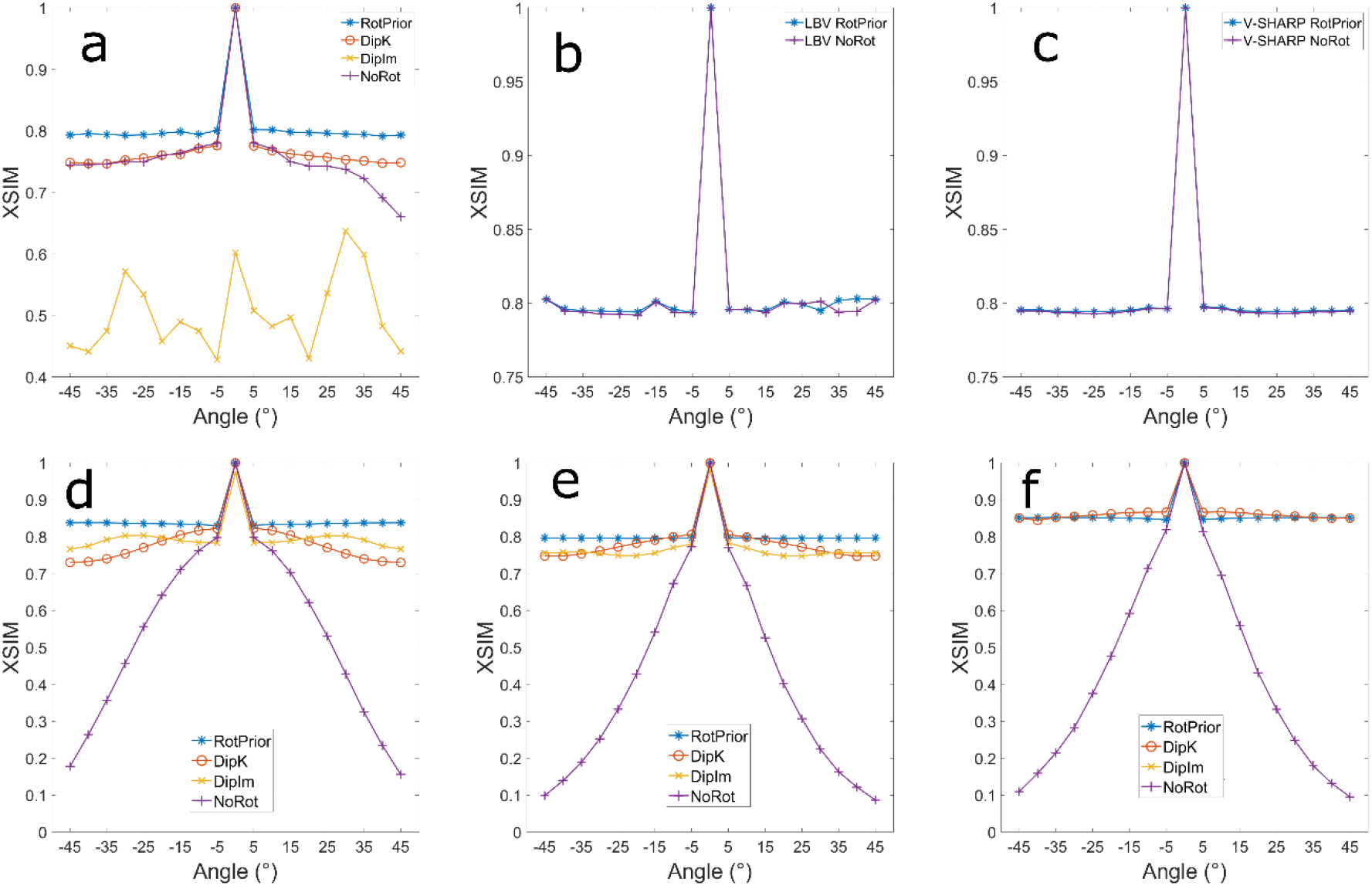
Results in the numerical phantom from oblique image volumes tilted about the x-axis. The volumes were padded to ensure no parts of the original image volume were cut off during rotations. XSIM measurements of QSM calculated with tilt corrections prior to background field removal with PDF (a), LBV (b) and V-SHARP (c) agree with unpadded results (Figure 5). XSIM measurements comparing tilt correction schemes prior to susceptibility calculation with TKD (d), iterative Tikhonov regularisation (e) and linear weighted linear TV (f) methods are also in agreement with unpadded results (Figure 6, bottom row).

**Supporting Figure S2:**
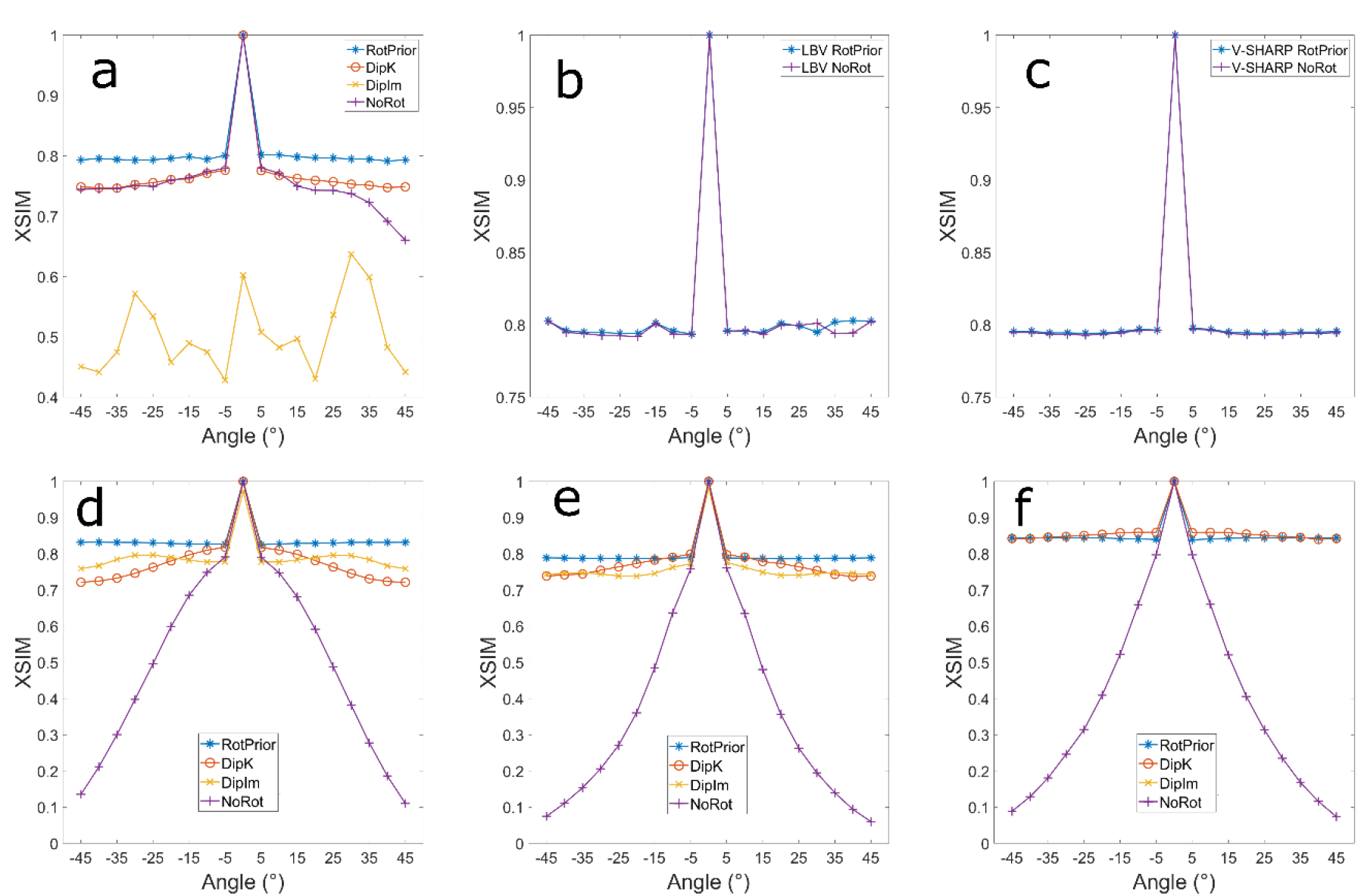
Results in the numerical phantom from oblique image volumes tilted about the *y*-axis with padded image volumes to ensure no parts of the original image volume were cut off during rotations. XSIM measurements of QSMs calculated with tilt corrections prior to background field removal with PDF (a), LBV (b) and V-SHARP (c) agree with unpadded results and rotations about the x-axis (Figure 5, Supporting Figure 1a,b,c). XSIM measurements comparing tilt correction schemes prior to susceptibility calculation with TKD (d), iterative Tikhonov regularisation (e) and linear weighted linear TV (f) methods are also in agreement with unpadded results and x-axis rotations (Figure 6, bottom row; Supporting figure 1 d, e, f).

**Supporting Figure S3:**
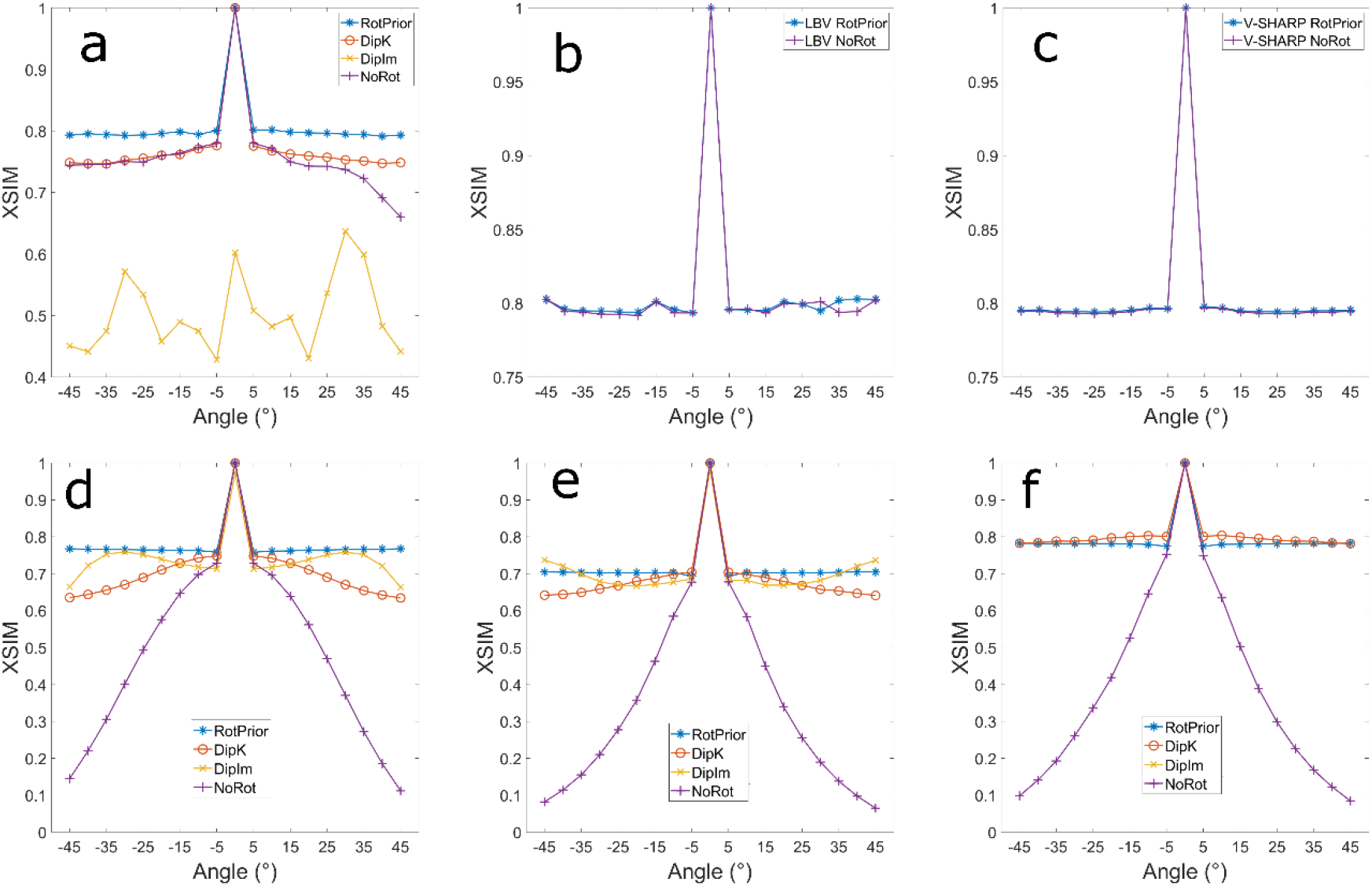
numerical phantom results from rotations about the y=x axis with padded image volumes to ensure no parts of the original image volume was cut off during rotations. XSIM measurements of QSMs during the background field removal part of the pipeline for PDF (a), LBV (b) and V-SHARP (c) agree with unpadded results and rotations about the x and y-axes. XSIM measurements comparing susceptibility calculation methods TKD (d), iterative Tikhonov regularisation (e) and linear weighted linear TV (f) are also in agreement with unpadded results and x and y axis rotations (Figure 6, bottom row; supporting figures 1 and 2 d, e, f).

**Supporting Figure S4:**
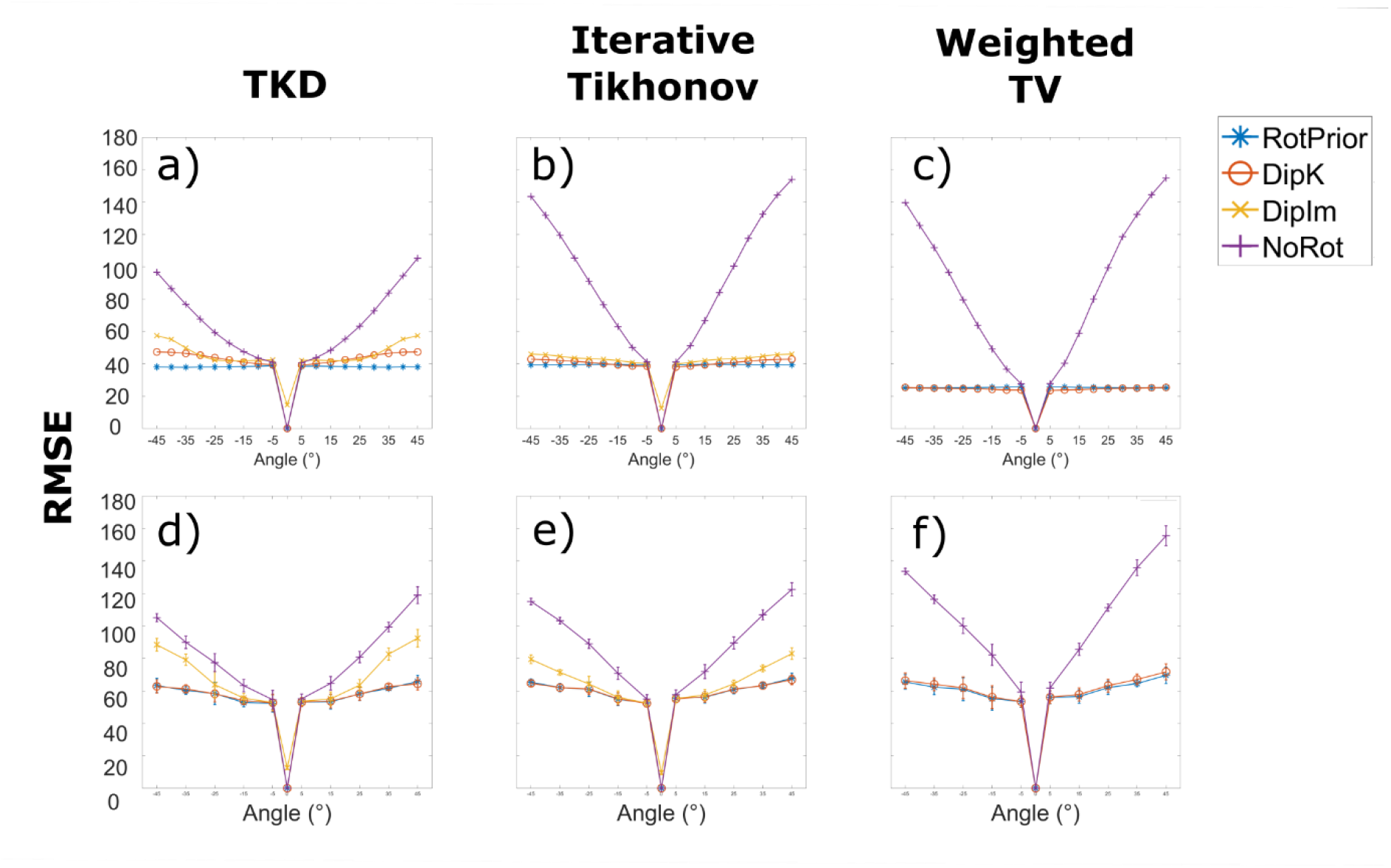
RMSE plots over all angles for all tilt correction schemes and all three susceptibility calculation methods in the numerical phantom (a-c) and averaged across all healthy volunteers (d-f). These results agree with the XSIM measurements found in Figures 6 and 9, for the numerical phantom and the in vivo results, respectively. Error bars (d-f) represent the standard deviation on the mean across all volunteers.

**Supporting Figure S5:**
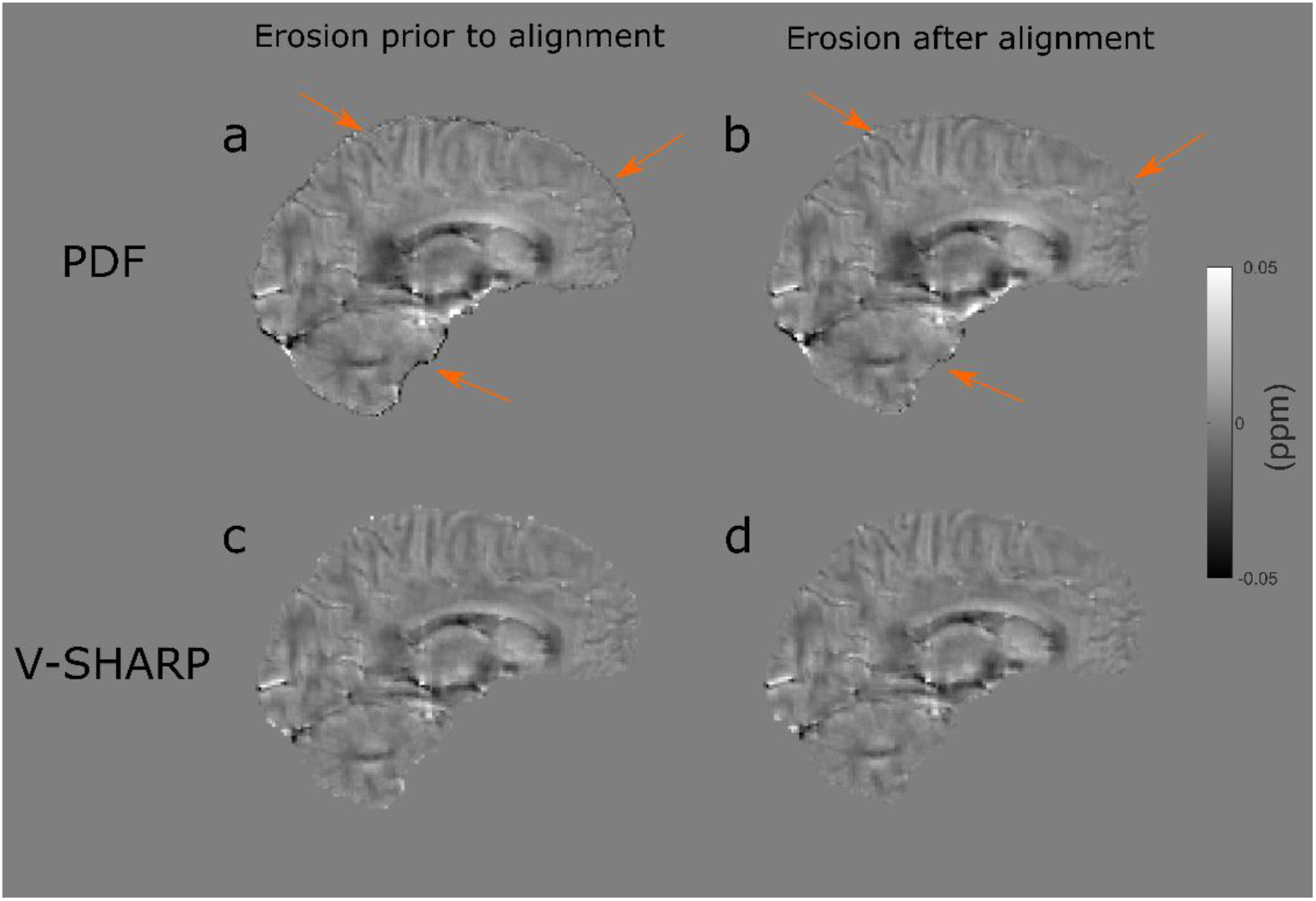
For improved PDF performance, the brain mask is typically eroded. If this erosion takes place prior to rotating the field map into alignment with 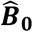 (a) compared to after (b), artefacts arise along the edges of the local field map following background field removal with PDF (a, orange arrows), increasing the RMSE and decreasing the XSIM. These artefacts do not arise when using V-SHARP (c: erosion before, d: erosion after).

**Supporting Figure S6:**
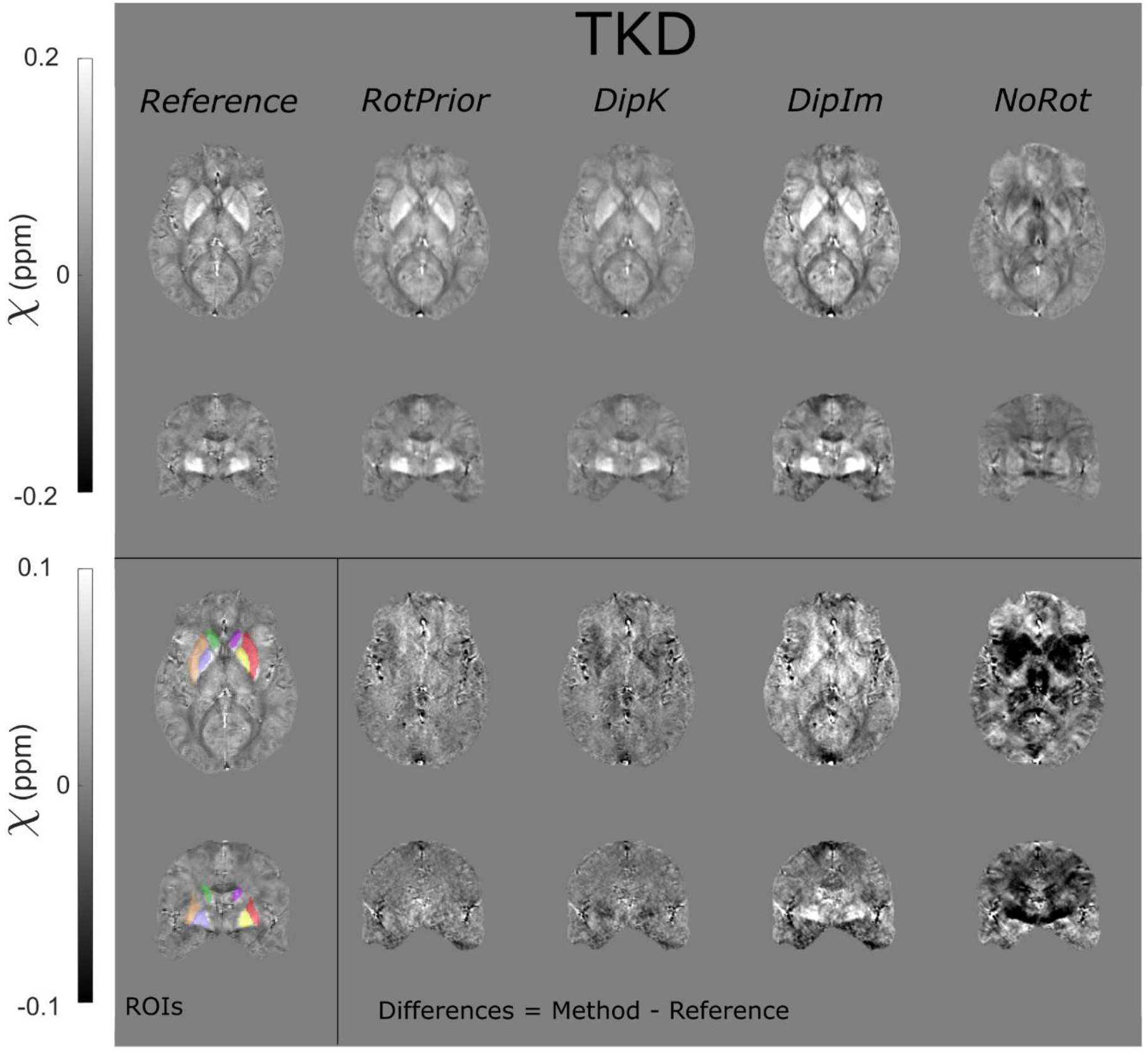
χ maps and difference images illustrating the effects of all tilt correction schemes on susceptibility calculation in vivo. An axial and a coronal slice are shown for a volume tilted at 45° and a reference (0°) volume with all χ maps calculated using Thresholded k-space (TKD) method. *NoRot* leads to the largest differences and image artefacts throughout the brain. The EVE ROIs used are shown (bottom left). These results are very similar to iterative Tikhonov and weighted linear TV susceptibility maps (Figure 10).

**Supporting Figure S7:**
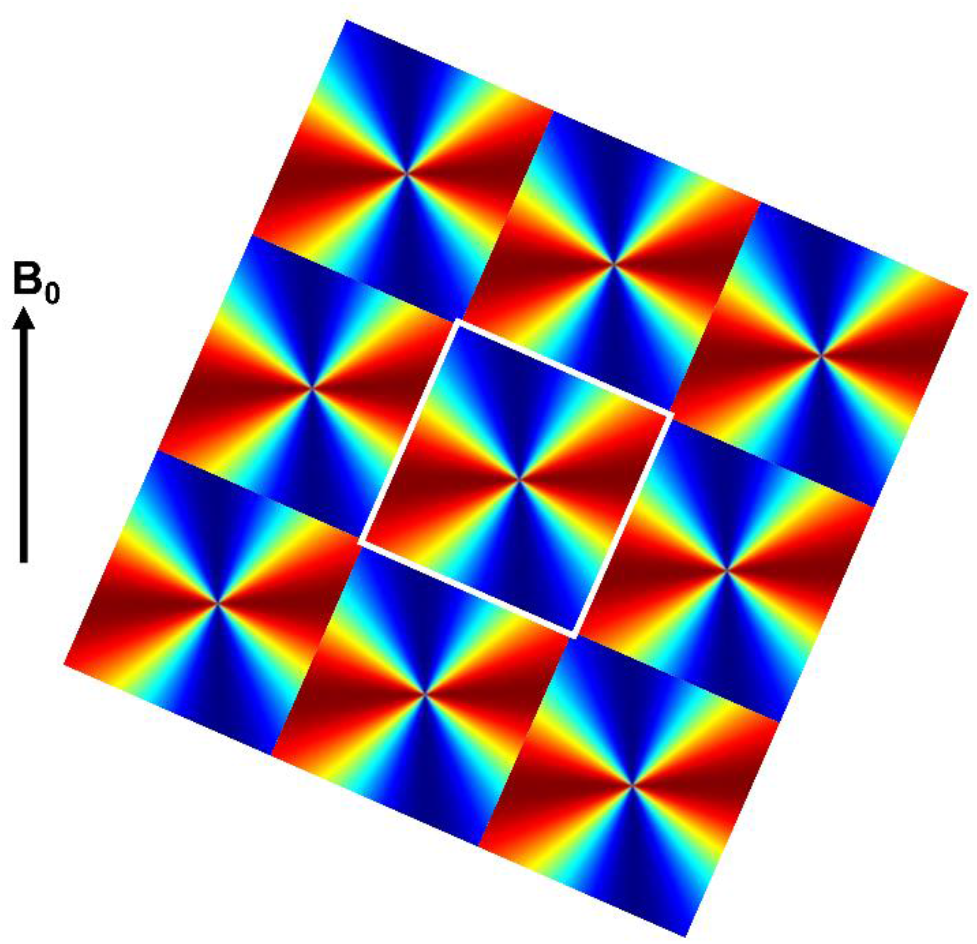
Oblique *k*-space magnetic dipole kernels laid side by side to illustrate the violations in circular continuity. These dipoles are used in the *DipK* correction method, which leads to striping artifacts due to the violations in circular continuity i.e. discontinuities at the boundaries of the rotated k-space dipoles (white square and arrows).

**Supporting Figure S8:**
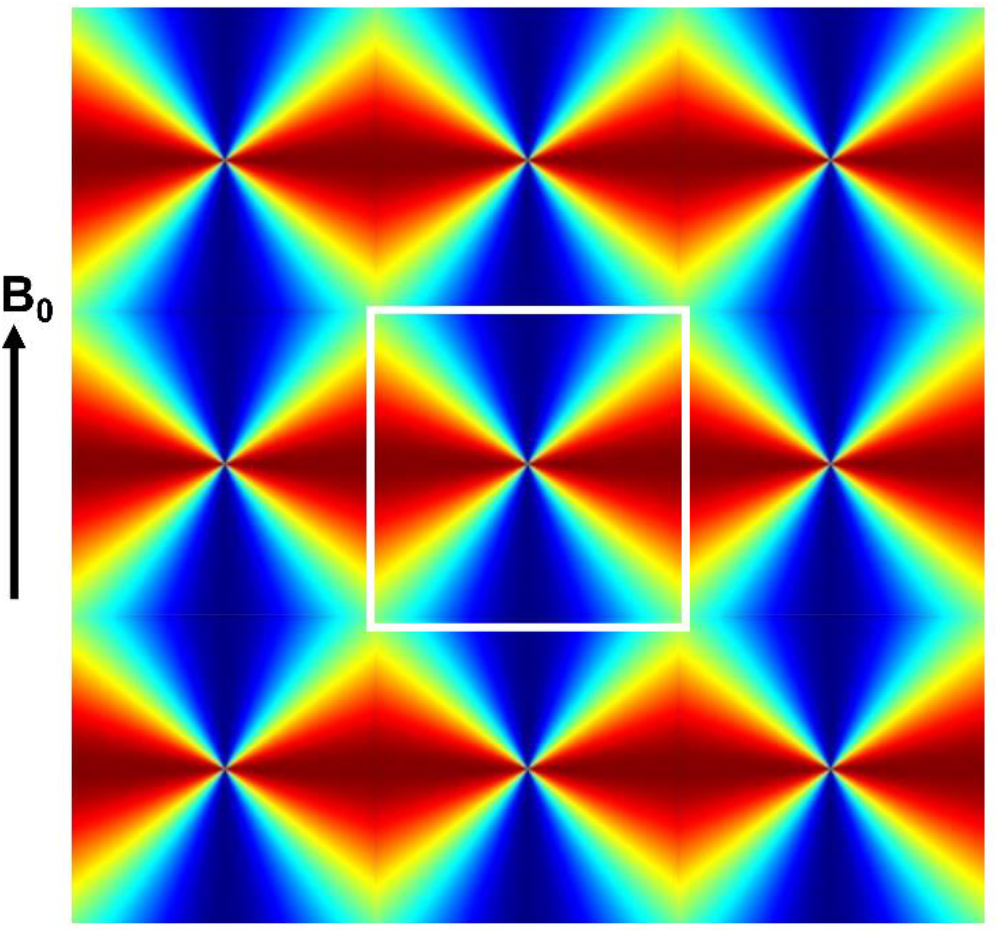
Non-oblique *k*-space magnetic dipole kernels laid side by side to illustrate circular continuity. When there is no oblique acquisition then there are no violations in circular continuity i.e. identical values and no discontinuities at the boundaries of the *k*-space dipoles (white square).

